# Centriole and PCM cooperatively recruit CEP192 to spindle poles to promote bipolar spindle assembly

**DOI:** 10.1101/2020.11.10.377341

**Authors:** Takumi Chinen, Kaho Yamazaki, Kaho Hashimoto, Ken Fujii, Koki Watanabe, Yutaka Takeda, Shohei Yamamoto, Yuka Nozaki, Yuki Tsuchiya, Daisuke Takao, Daiju Kitagawa

## Abstract

The pericentriolar material (PCM) that accumulates around the centriole expands during mitosis and nucleates microtubules. While centrosomes facilitate bipolar spindle formation, the individual functions of the centriole and PCM in mitosis remain elusive. Herein, we show the redundant roles of the centriole and PCM in bipolar spindle formation in human cells. Upon depletion of the PCM scaffold components, pericentrin and CDK5RAP2, centrioles remained able to recruit CEP192 onto their walls, which was sufficient for bipolar spindle formation. In contrast, through centriole removal, we found that pericentrin and CDK5RAP2 recruited CEP192 at the acentriolar spindle pole and facilitated bipolar spindle formation in mitotic cells with one centrosome. Furthermore, the chemical perturbation of polo-like kinase 1, a critical kinase for PCM assembly, efficiently suppressed the proliferation of various cancer cell lines from which centrioles were removed. Overall, these data suggest that the centriole and PCM cooperatively recruit CEP192 to spindle poles and facilitate bipolar spindle formation in human cells.

## Introduction

Centrosomes nucleate and anchor microtubules, thereby facilitating efficient spindle formation and chromosome segregation during mitosis(Moritz et al., 1995; Kollman et al., 2011; Woodruff et al., 2017). The microtubule-organizing activity of centrosomes depends on the pericentriolar material (PCM) that surrounds one or two centrioles(Hyman, 2014). Abnormalities in centrosome organization and function lead to chromosomal segregation errors; several mutations in centrosomal proteins have also been implicated in the development of diseases such as cancer(Nigg and Raff, 2009; Gönczy, 2015). In addition, PCM disorganization directly causes chromosome mis-segregation(Watanabe et al., 2019; Cosenza et al., 2017). Therefore, elucidating the function and organization of centrosome in mitosis will contribute to a better understanding of the mechanisms through which centrosomes dictate the spindle structure and support accurate chromosome segregation.

PCM contains a large number of proteins, such as the γ-tubulin ring complex (γ-TuRC), CDK5RAP2, CEP192, and pericentrin. During the G2/M transition, CEP192 recruits Aurora A and polo-like kinase 1 (PLK1) to centrosomes in a pericentrin-dependent manner; subsequently, CEP192 activates these kinases to promote microtubule nucleation and centrosome separation(Joukov et al., 2014). CEP192 also supports the organization of other PCM components for efficient bipolar spindle assembly(Gomez-Ferreria et al., 2007). PLK1 phosphorylates pericentrin to further recruit other PCM components to centrosomes, thereby increasing the microtubule nucleation activity of the centrosome during mitosis(Lee and Rhee, 2011). Microtubule nucleation activity depends on γ-TuRC(Zheng et al., 1995; Wieczorek et al., 2020; Liu et al., 2020; Consolati et al., 2020; Moritz et al., 1995; Kollman et al., 2011), the activity of which is upregulated by the binding of CDK5RAP2 to γ-TuRC(Choi et al., 2010; Hanafusa et al., 2015). In addition to their functions in microtubule nucleation, previous studies have described pericentrin and CDK5RAP2 regulating spindle pole focusing and spindle orientation through the regulation of motor proteins or other spindle pole proteins(Lee and Rhee, 2010; Chavali et al., 2016; Tungadi et al., 2017; Chen et al., 2014).

During the G2/M phase, PCM expands around the pair of centrioles that form the structural core of the centrosome, and increases its ability to nucleate microtubules. In *Drosophila* and *Caenorhabditis elegans*, it has been reported that the centrioles regulate the architecture and dynamics of PCM(Kirkham et al., 2003; Conduit et al., 2010; Erpf et al., 2019; Cabral et al., 2019; Sir et al., 2013; Alvarez-Rodrigo et al., 2019; Conduit et al., 2014). In addition, it has been shown that PCM disorganization causes precocious centriole disengagement during mitosis(Seo et al., 2015; Kim et al., 2015, 2019; Watanabe et al., 2019), which can result in impairment of spindle pole integrity(Watanabe et al., 2019). This cross-reactive interplay between centrioles and PCM complicates the analysis of the individual function of PCM at spindle poles independent from the involvement of centriolar machinery. The centriole-independent functions of PCM have been partially characterized in the acentriolar meiotic spindles of mouse oocytes. During meiotic spindle formation in mice, acentriolar microtubule-organizing centers are formed and merge into two equal spindle poles(Clift and Schuh, 2015; Schuh and Ellenberg, 2007). Conditional knockout of pericentrin induces spindle instability and severe meiotic errors that lead to pronounced female subfertility in mouse oocytes. These findings suggest that pericentrin assists in organizing functional spindle poles to achieve faithful chromosome segregation(Baumann et al., 2017). However, as the system of meiosis is particularly unique compared with that of mitosis, it is unclear whether acentrosomal spindle formation pathways can be directly compared between oocytes and somatic cells.

To evaluate the distinct functions of PCM in human somatic cells independently of centrioles, it is important to utilize an assay system that enables the analysis of mitotic spindles that lack centrioles. As centriole duplication requires PLK4(Habedanck et al., 2005; Bettencourt-Dias et al., 2005), its specific inhibitor centrinone can be used to remove centrioles(Wong et al., 2015). Treatment with centrinone leads to progressive loss of centrioles and generates mitotic spindles with one or zero centrosomes. Using this strategy, we have previously shown the critical roles of NuMA in the spindle bipolarization in early mitosis of cells without centrosomes(Chinen et al., 2020). Similarly, by using mitotic cells with one centrosome, Dudka et al. recently reported that centrosomes regulate the length of K-fibers, and thereby alter their dynamics in HURP-dependent manner(Dudka et al., 2019).

In this study, we show the redundant roles of the centriole and PCM in bipolar spindle formation in human cells. When PCM assembly was inhibited by depletion of the PCM scaffold proteins pericentrin and CDK5RAP2, we found that CEP192 remained at the centriole wall, where it presumably promoted bipolar spindle formation. Furthermore, we induced the formation of mitotic spindles with only one centrosome by treating human cells with centrinone. We found that the one-centrosome cells formed a bipolar spindle that accumulated PCM components, including CEP192, at the acentriolar pole. In such cells, depletion of pericentrin or CDK5RAP2 compromised the formation of the acentriolar pole and significantly prolonged mitotic progression. In contrast, the artificial accumulation of PCM components at the acentriolar pole accelerated the mitotic progression in one-centrosome cells. These results demonstrate that the centriole and PCM cooperatively assemble CEP192 at the spindle poles and facilitate bipolar spindle formation in human cells.

## Results

### CEP192 at the centriolar wall is sufficient for organizing mitotic spindle poles

To understand the functions of PCM in bipolar spindle formation in human cells, we depleted the main components of PCM, such as CEP192, pericentrin, and CDK5RAP2, and observed mitotic progression in HeLa cells. As previously described, the depletion of CEP192 caused severe defects in bipolar spindle formation and prolonged mitotic duration (Fig. 1A, B). On the other hand, double depletion of pericentrin and CDK5RAP2 or their individual depletion had a limited effect on mitotic duration (Fig. 1A, B, 2C, and S1G). These results suggest that CEP192, but not pericentrin or CDK5RAP2, is critical for mitotic progression. It has been suggested that pericentrin and CDK5RAP2 cooperatively recruit PCM components, including CEP192, at centrosomes(Kim and Rhee, 2014). Therefore, we observed the amount and localization of CEP192 at centrosomes upon depletion of pericentrin and CDK5RAP2. We found that a certain quantity of CEP192 remained at centrosomes in pericentrin/CDK5RAP2 double-depleted cells (Fig. 1C and D). To further understand this mechanism, we used gated stimulated emission depletion (STED) microscopy to analyze the detailed localization pattern of CEP192 at centrosomes in pericentrin/CDK5RAP2 double-depleted cells (Fig. 1E–G). Centrioles were marked by poly-glutamylated centriolar microtubules. In control cells, CEP192 was detectable in the PCM clouds that surrounded mother centrioles (Fig. 1E–G). In contrast, in pericentrin/CDK5RAP2 double-depleted cells, the reduced quantity of CEP192 was detectable only on centriole walls. These results raise the possibility that CEP192 at the centriolar wall, rather than in the PCM cloud, is crucial for the microtubule-organizing center function of centrosomes.

**Figure 1.**
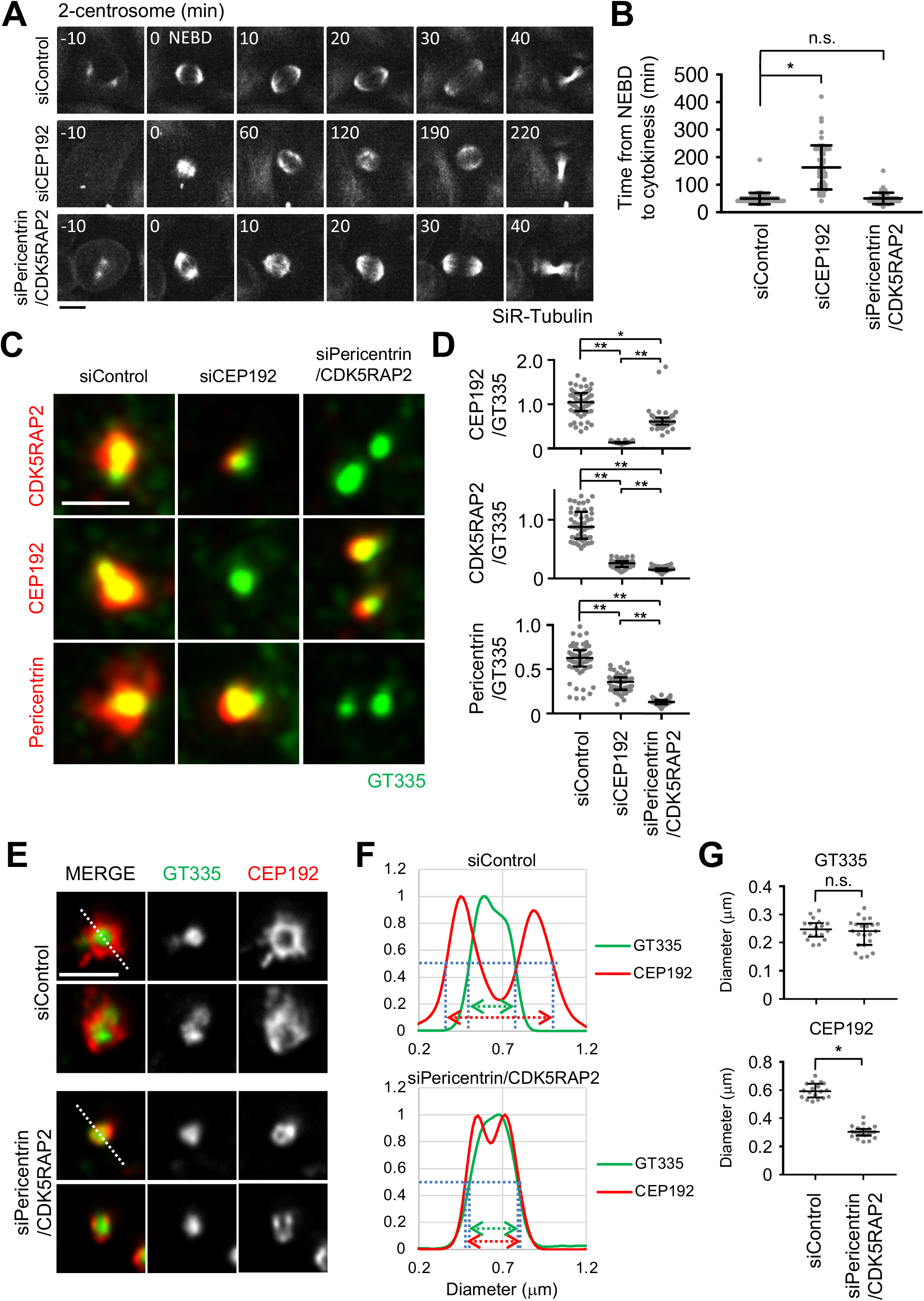
CEP192 at centriolar wall, but not the expanded PCM mediated by pericentrin and CDK5RAP2, is sufficient for bipolar spindle formation. (**A**) Time-lapse observation of the structure of microtubules upon siRNA treatment against the indicated proteins. HeLa cells expressing EGFP-centrin1 and pericentrin-mCherry were observed with a 40× objective. Gray represent SiR-tubulin, respectively. Z-projections: 10 planes, 2.2 µm apart. Scale bar, 10 µm. Time zero corresponds to nuclear envelope break down (NEBD). (B) Mitotic duration, the time required from NEBD to cytokinesis, in (A). Line and error bars represent the mean and SD (N ≥ 50 cells from two independent experiments). Kruskal–Wallis test was used to determine the significance of the difference. *P < 0.05. (**C**) The localization of PCM proteins in mitotic spindles of the cells in which the indicated protein was depleted. Red and green represent PCM proteins (CDK5RAP2, Cep192 or PCNT) and GT335, respectively. Z-projections of 10 sections, every 0.3 μm. Scale bar, 1 μm. (**D**) The signal intensity of PCM proteins on mitotic centrosomes of fixed HeLa cells was analyzed (N>45 for each condition). Line and error bars represent median with interquartile range. Kruskal–Wallis test was used to determine the significance of the difference. *P < 0.05, **P < 0.0001. (**E**) STED images showing centriolar distribution of Cep192 in PCNT/CDK5RAP2 double-depleted cells. HeLa cells were treated with control siRNA or PCNT/CDK5RAP2 siRNA for 48 h and stained with the indicated antibodies. Scale bar, 1 μm. (**F, G**) Representative line intensity profiles (F) and measured diameters (G) of GT335 and Cep192. The line profiles were measured along the dotted lines in (E). The profiles were fitted with double Gaussian curves and the distances between the half-maximal intensity points at the far ends were measured as the diameters (schematically indicated with dotted lines and arrows in the profiles; fitted curves are not shown). Horizontal bars and error bars in the plots for the diameters represent median and interquartile range. *N* = 18 (for siControl) or 22 (for siPCNT/CDK5RAP2) centrosomes; data from two independent experiments were pooled. Mann–Whitney U-test was used to determine the significance of the difference. *P < 0.0001.

**Figure 2.**
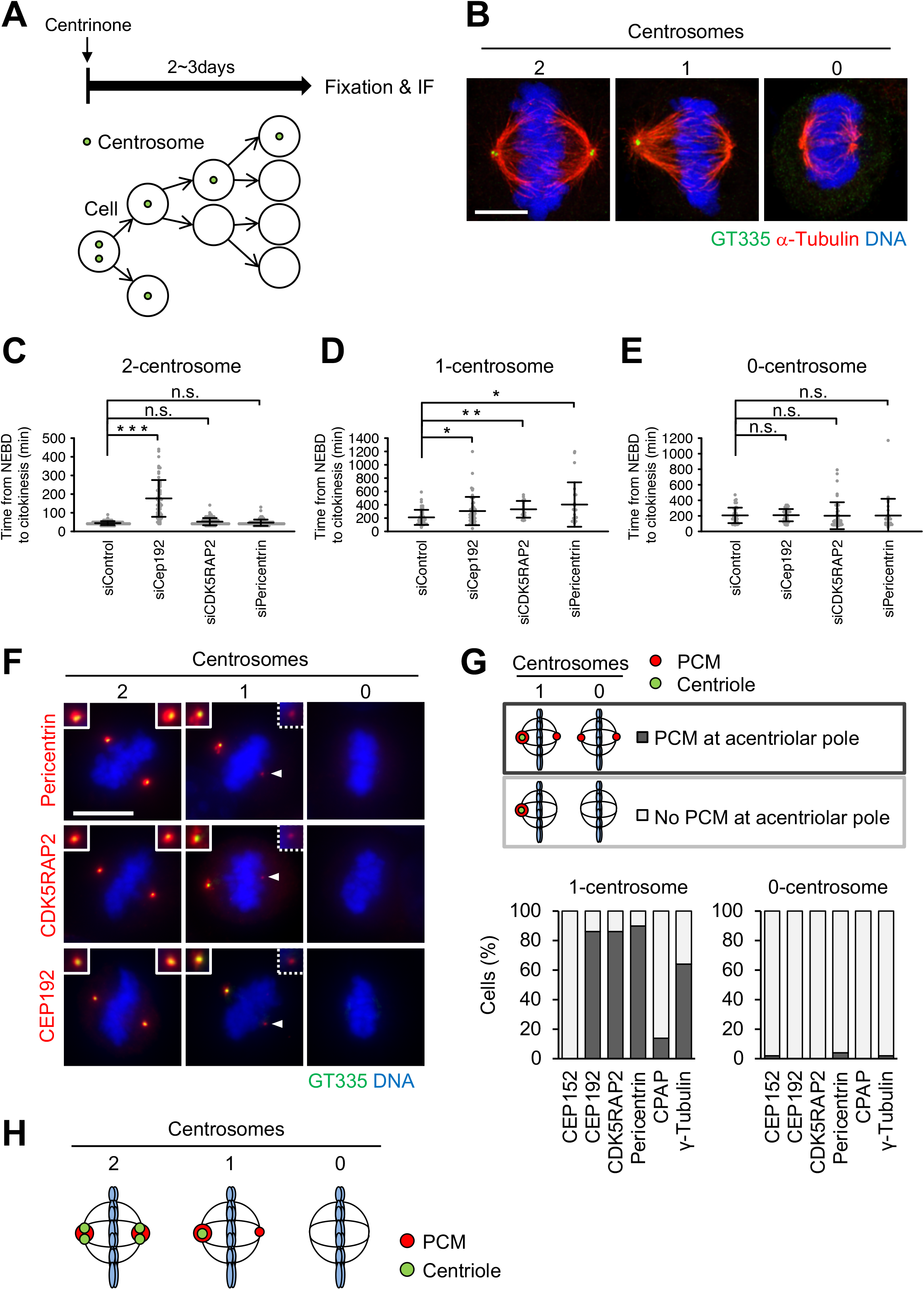
Cells with one centrosome can organize bipolar spindles in mitosis by forming a PCM-positive acentriolar spindle pole (PCM-pole). (**A**) Schematic illustration of centrinone-induced removal of centriole. (**B**) DMSO-treated control mitotic spindles (two centrosomes) and centrinone B-treated centrosome-depleted spindles (one or zero centrosomes). Green, red, and blue represent GT335 (polyglutamylated centriole microtubules), α-tubulin, and DNA, respectively. Z-projections: 12 planes, each 0.13 μm apart. Scale bar, 5 μm. (**C-E**) Mitotic duration, the time required from NEBD to cytokinesis, in DMSO-treated two-centrosome (C), centrinone-treated one-centrosome (D) and zero-centrosome (E) cells, in Fig (S1G-I). Line and error bars represent the mean and SD (N ≥ 20 cells from two independent experiments). Kruskal–Wallis test was used to determine the significance of the difference. \**P* < 0.05, \*\**P* < 0.005, *** *P* < 0.0001. (**F**) Distribution of centrosomal factors in centriolar and acentriolar spindle poles. DMSO-treated control mitotic spindles (two centrosomes) and centrinone-treated mitotic spindles (one or zero centrosome) in HeLa cells. Green, red, and blue represent GT335, the protein of interest (Pericentrin, CDK5RAP2, or CEP192), and DNA, respectively. Z-projections: 21 sections, every 1μm. Scale bar, 10 μm. (**G**) Quantification of pole patterns in (F). Values are presented as mean percentages from two independent experiments (N = 25 for each experiment). (**H**) Schematic illustration of PCM localization at spindle poles in two-, one- or zero-centrosome cells.

### Cells with one centrosome form a bipolar spindle that accumulates PCM components at the acentriolar pole

To understand the functions of PCM independently of centrioles in human cells, we next induced the formation of mitotic spindles with one or zero centrosomes by treating HeLa cells with the PLK4 inhibitors centrinone or centrinone B (Fig. 2A and B). Centrosomes were marked by polyglutamylated centriolar microtubules or centrin to determine their number. We depleted PCM components CEP192, pericentrin, and CDK5RAP2 in one- or zero-centrosome cells, and observed their mitotic progression using live cell imaging. As described above, the depletion of CEP192, but not pericentrin or CDK5RAP2, prolonged mitosis in cells with two centrosomes (Figs. 2C, S1A–G). On the other hand, interestingly, we found that depletion of pericentrin or CDK5RAP2, as well as CEP192, significantly prolonged mitotic duration in one-centrosome cells (Figs. 2D, S1H). In contrast, we found that depletion of pericentrin, CDK5RAP2, or CEP192 had a limited effect on mitotic progression in zero-centrosome cells (Figs. 2E, S1I). These results suggest that pericentrin and CDK5RAP2 are important for mitotic progression in one-centrosome cells, but not in two- or zero-centrosome cells.

We further analyzed the localization patterns of PCM proteins at spindle poles using immunofluorescence microscopy (Figs. 2F, G, S2A–D). We found that the acentriolar spindle poles of one-centrosome cells incorporate a detectable amount of PCM components, such as pericentrin, CDK5RAP2, CEP192, and γ-tubulin (Figs. 2F, G, S2A) but not CEP152 or CPAP (Figs. 2G and S2A). In this study, we termed the acentriolar spindle pole that contains PCM the ‘PCM-pole’. In contrast, most spindle poles of zero-centrosome cells lacked PCM components, as previously described (Figs. 2F, G, S2A) (Chinen et al., 2020). PCM components were consistently detectable at the acentriolar spindle poles in one-centrosome cells of various human cell lines (Fig. S2B). Furthermore, the PCM-pole was similarly observed in one-centrosome cells induced by SAS6-depletion using the auxin-inducible degron (AID) system (Fig. S2C, D)(Yoshiba et al., 2019), suggesting that this phenotype was not a specific result of PLK4 inhibition.

We next examined whether the PCM-pole nucleates microtubules using a microtubule regrowth analysis. For this analysis, we immunostained the microtubule end binding protein 1 (EB1), which marks growing microtubule plus ends. When restarting the microtubule nucleation, the EB1 signals started developing around both centriolar and PCM-poles, with PCM-poles nucleating less microtubules (Fig. S2E). Thus, PCM-poles possess microtubule nucleation activity, although this activity appears slightly lower than that of centriolar poles. Collectively, these results suggest that one-centrosome cells assemble PCM at the acentriolar spindle pole, which harbors microtubule nucleation activity (Fig. 2H).

### The PCM-pole is formed by either split of the PCM from the centriolar pole or accumulation of PCM

To understand the mechanism of PCM recruitment to the acentriolar pole in one-centrosome cells, we used time-lapse fluorescence microscopy to track the dynamics of endogenous pericentrin or CDK5RAP2 tagged with mCherry as markers of PCM. This strategy revealed that, at first, pericentrin accumulated at centriolar poles in early mitosis. Subsequently, one-centrosome cells formed pericentrin-positive PCM-poles by either splitting of the PCM from the centriolar pole or *de novo* accumulation of PCM (38.5% and 51.9%, respectively) (Fig. 3A, B, D). These PCM-poles disappeared after cytokinesis (Fig. 3A, B, E), consistent with the observation that PCM proteins are disassembled after mitotic exit(Woodruff et al., 2014). On the other hand, a detectable amount of pericentrin did not accumulate at the acentriolar spindle poles in most zero-centrosome cells (Fig. 3C, D). Taken together, these observations suggest that one-centrosome cells initially accumulate PCM proteins around centrioles and subsequently generate the acentriolar pole by splitting and/or by recruiting PCM components on the opposite side for bipolar spindle formation (Fig. 3F).

**Figure 3.**
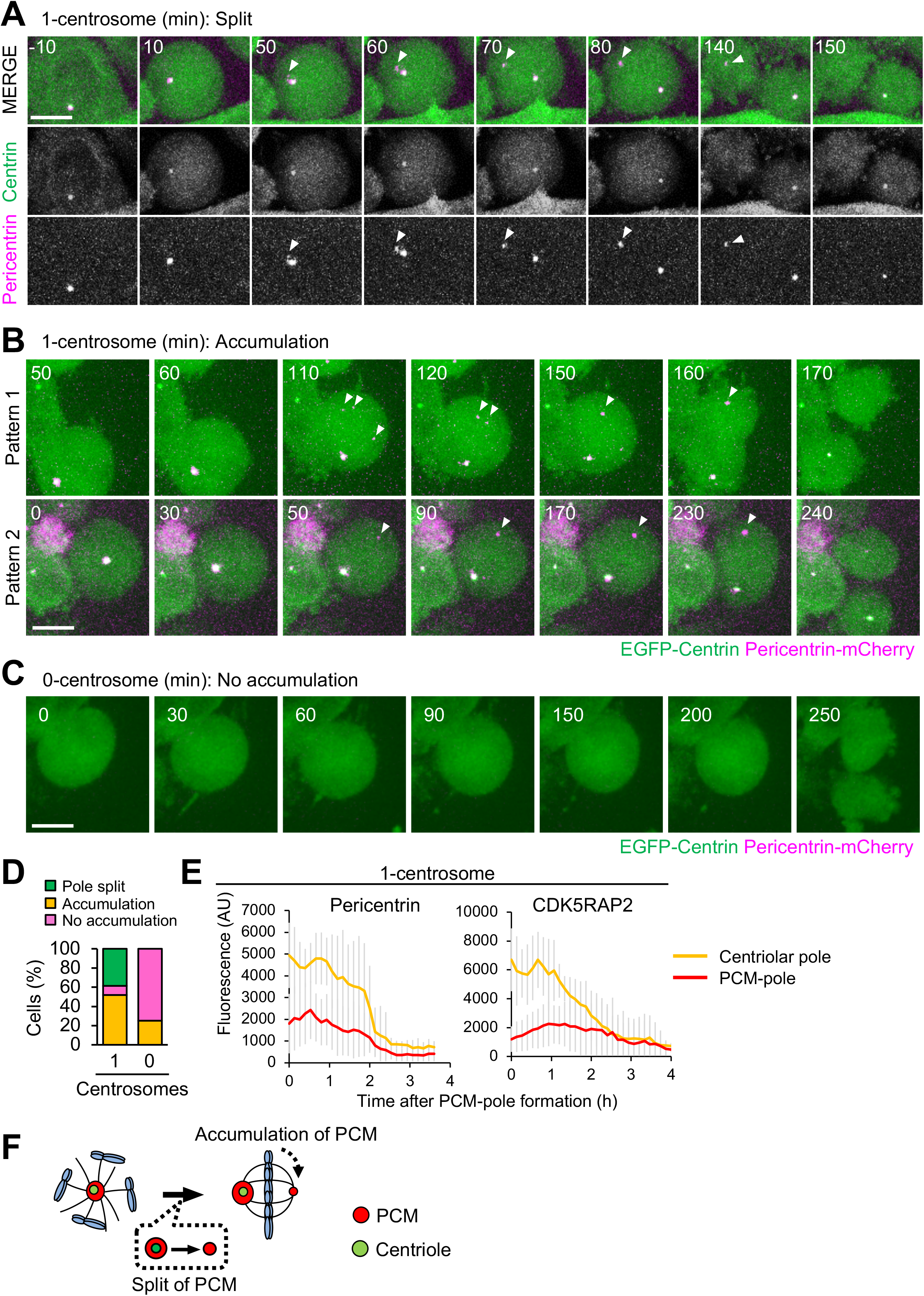
The PCM-pole is formed by splitting PCM from the centriolar pole or by accumulation of PCM components. (**A–D**) HeLa cells expressing EGFP-centrin1 and pericentrin-mCherry were observed with a 60× objective. Magenta and green represent pericentrin and centrin, respectively. Z projections: 20 planes, 1.2 μm apart. Scale bar, 10 μm. Time zero corresponds to the beginning of mitotic cell rounding. (**A**) Splitting of the PCM components from the centriolar pole in one-centrosome cells. Arrowheads indicate the PCM at the acentriolar spindle pole. (**B**) PCM accumulation in one-centrosome cells. Arrowheads indicate the accumulation of PCM at acentriolar spindle poles. (**C**) Cell division in zero-centrosome cells without accumulation of PCM. (**D**) Quantification of patterns of PCM dynamics in (A–C). Values are percentages of the total cells from 52 (for one-centrosome cells) or 24 (for zero-centrosome cells) cells from two independent experiments. (**E**) Averaged time courses of pericentrin-mCherry or CDK5RAP2-mCherry signals at the centriolar spindle pole and PCM-pole of 10 cells. Time course data were aligned at PCM-pole formation (0 h). Error bars, SD; A.U., arbitrary units. (**F**) Schematic illustration of PCM-pole formation by splitting PCM from the centriolar spindle pole or by accumulation of PCM components.

### CDK5RAP2 and pericentrin are crucial for the bipolar spindle formation in one-centrosome cells

We next analyzed the specific role of PCM in cell division in one-centrosome cells. However, it is difficult to analyze the specific roles of some PCM-pole components. Among those appeared to localize at PCM-poles (Fig. 2F, G, S2A), for example, γ-tubulin also localizes along the whole spindle and regulates several pathways of microtubule nucleation in mitosis(Lecland and Lüders, 2014; Teixido-Travesa et al., 2012). In addition, CEP192 is required for bipolar spindle formation in cells with two centrosomes(Zhu et al., 2008; Joukov et al., 2014). On the other hand, depletion of the PCM scaffold proteins CDK5RAP2 and pericentrin are known to have little effect on spindle formation in two-or zero-centrosome cells (Fig. 2C-E, S1G-I). Therefore, we selected CDK5RAP2 and pericentrin for further analysis of PCM-poles in one-centrosome cells.

We found that depletion of CDK5RAP2 or pericentrin caused arrest of one-centrosome cells in mitosis with monopolar spindles; however, this effect was not observed in two-centrosome cells (Fig. 4A, B). These results indicate that CDK5RAP2 and pericentrin play an important role in bipolar spindle formation specifically in one-centrosome cells, but not in two-centrosome cells. To further investigate this process, we tracked the dynamics of spindle poles in one-centrosome cells using time-lapse observation of NuMA tagged with mCherry. Upon depletion of CDK5RAP2 or pericentrin, the separation of two NuMA foci was normally detectable in early mitosis (Fig. 4C, D), while the time from nuclear envelope breakdown to cytokinesis was prolonged (Fig. 4C–F). In addition, immunofluorescence analysis revealed that, in the pericentrin or CDK5RAP2-depleted HeLa cells with one-centrosome, the degree of CEP192 localization at the acentrosomal spindle poles was reduced (Fig. 4G). These results indicate that PCM scaffold proteins CDK5RAP2 and pericentrin are crucial for the recruitment of CEP192 at the acentrosomal spindle pole and bipolar spindle formation, but are likely dispensable for the early step of spindle pole generation in one-centrosome cells.

**Figure 4.**
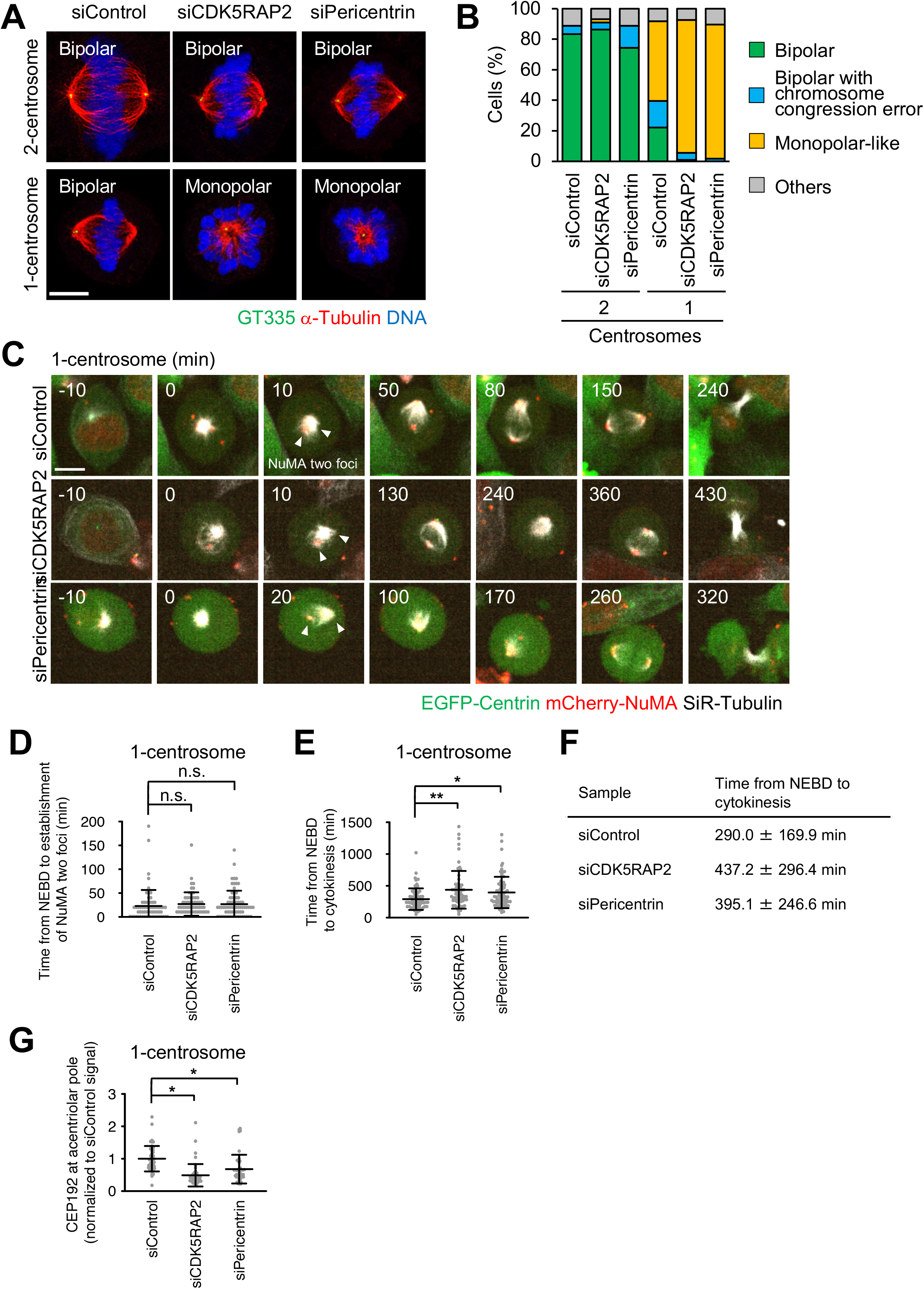
Pericentrin and CDK5RAP2 are crucial for spindle elongation and spindle bipolarization of one-centrosome cells. (**A**) Mitotic spindle structures upon siRNA treatment with or without 500 nM of CentB. Green, red, and blue represent GT335, α-tubulin, and DNA, respectively. Z-projections: 5 planes, 0.3 μm apart. Scale bar, 5 μm. (**B**) Frequency of mitotic spindle structures after siRNA treatment against the indicated proteins in (A). Values are presented as mean percentages. N > 86, data from two independent experiments were pooled. (**C**) Time-lapse observation of the structure of NuMA and microtubules upon siRNA treatment against the indicated proteins. Centrinone-treated one-centrosome HeLa cells expressing EGFP-centrin1 and pericentrin-mCherry were observed with a 40× objective. Red, green, and gray represent NuMA, centrin, and, SiR-tubulin, respectively. Z-projections: 10 planes, 2.2 μm apart. Scale bar, 10 μm. Time zero corresponds to nuclear envelope break down (NEBD). Arrowheads indicate the separated two NuMA foci. (**D**) The time required for the initial establishment of two poles of NuMA in (C). Line and error bars represent the mean and SD (N ≥ 60 cells from three independent experiments). Kruskal–Wallis test was used to determine the significance of the difference. n.s., not significantly different (*P* > 0.05). (**E**) Mitotic duration, the time required from NEBD to cytokinesis, in (C). Line and error bars represent the mean and SD (N ≥ 60 cells from three independent experiments). Kruskal–Wallis test was used to determine the significance of the difference. \**P* < 0.01, \*\**P* < 0.001. (**F**) Table of the times from nuclear envelope break down (NEBD) to cytokinesis in (E). (**G**) The signal intensity of CEP192 on acentrosomal spindle poles. Line and error bars represent the mean and SD (N ≥ 46 cells from two independent experiments). Kruskal–Wallis test was used to determine the significance of the difference. *P < 0.0001.

To verify whether pericentrin and CDK5RAP2 are important for bipolar spindle formation in other human cell lines with one centrosome, we observed the spindle structure of RPE1 and A549 cells upon depletion of pericentrin or CDK5RAP2. Through immunostaining, we found that in cells with one centrosome, depletion of pericentrin or CDK5RAP2 induced the formation of monopolar spindles (Fig. S3A–D). These results further support the conclusion that PCM proteins are required for bipolar spindle formation in one-centrosome cells.

### Depletion of CEP57 promotes accumulation of PCM components at PCM-poles and facilitates bipolar spindle formation in one-centrosome cells

Next, we sought to further analyze the importance of PCM components at PCM-poles for cell division in one-centrosome cells. Since siRNA-mediated depletion reduces the total expression level of CDK5RAP2 and pericentrin, it is difficult to analyze the function of the PCM components specifically at PCM-poles (Fig. 4). Therefore, we used another approach to indirectly manipulate the amount of PCM components at PCM-poles: depleting CEP57. CEP57 provides a critical interface between the centriole and PCM, and depletion of CEP57 induces the fragmentation of PCM proteins in early mitosis of human cells(Watanabe et al., 2019). Given that 38.5% of one-centrosome cells assembled PCM-poles by splitting PCM from the centrosome (Fig. 3A, D), we hypothesized that, upon CEP57 depletion, the PCM that is dissociated from the centrosome could be incorporated into the acentriolar pole in one-centrosome cells. As expected, the amount of pericentrin at PCM-poles was significantly increased, presumably due to the increased PCM fragmentation at centriolar poles after CEP57 depletion (Fig. 5A–C). Subsequently, to analyze the effect of CEP57 depletion on the mitotic processes of one-centrosome cells, we performed time-lapse imaging of NuMA and microtubules. We found that depletion of CEP57 promoted bipolar spindle formation more efficiently than in control cells, and thereby shortened the mitotic duration (Fig. 5D–F). Under this condition, CEP57-depleted cells with one centrosome formed two separate NuMA foci, similar to siControl-treated one-centrosome cells, but established a bipolar spindle formation more efficiently (Fig. 5E, G). Overall, these results suggest that accumulation of PCM components at PCM-poles facilitates the bipolar spindle formation in one-centrosome cells.

**Figure 5.**
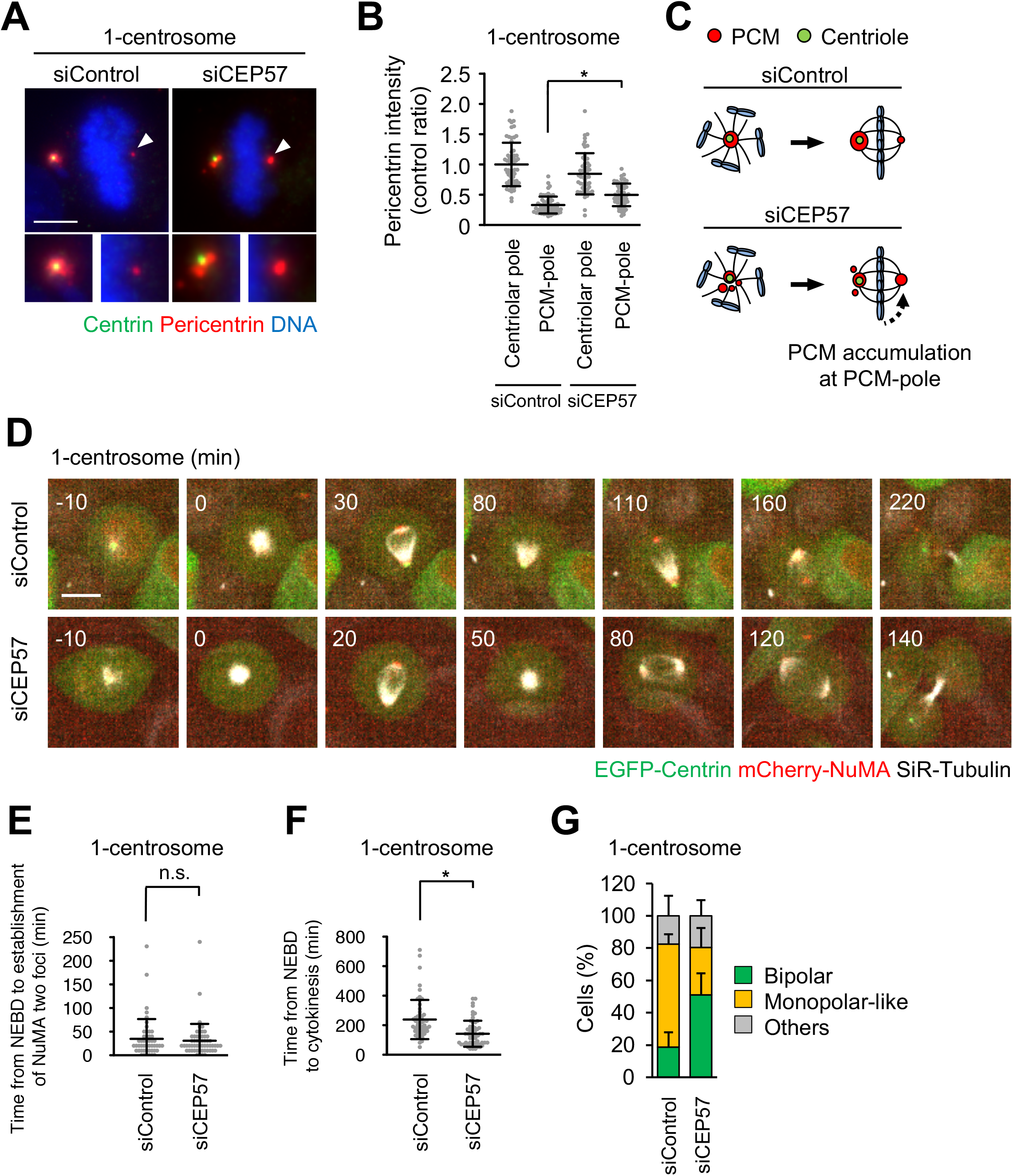
CEP57 depletion leads to an increase of PCM at the acentriolar pole and facilitates spindle bipolarization in one-centrosome cells. (**A**) Mitotic spindle pole structures of one-centrosome cells upon CEP57 depletion. Green, red, and blue represent centrin, pericentrin, and DNA, respectively. Z-projections: 20 planes, 0.5 μm apart. Scale bar, 5 μm. (**B**) The signal intensity of pericentrin on centrosomes or PCM-poles in (A). Line and error bars represent the mean and SD (N ≥ 50 cells from two independent experiments). Kruskal–Wallis test was used to determine the significance of the difference. \**P* < 0.01. (**C**) Schematic illustration of CEP57-depletion-induced pericentrin accumulation at the PCM-pole. (**D**) Time-lapse observation of NuMA structures and microtubules upon CEP57 depletion. Centrinone-treated one-centrosome HeLa cells expressing EGFP-centrin1 and pericentrin-mCherry were observed with a 40× objective. Red, green, and gray represent NuMA, centrin, and SiR-tubulin, respectively. Z-projections: 10 planes, 2.2 μm apart. Scale bar, 10 μm. Time zero corresponds to mitotic onset. (**E**) The time required for the initial establishment of two poles of NuMA in (D). Line and error bars represent the mean and SD (N ≥ 50 cells from two independent experiments). The Mann–Whitney *U*-test (two-tailed) was used to obtain a *P*-value. n.s., not significantly different (*P* > 0.05). (**F**) Mitotic duration, the time required from nuclear envelope break down (NEBD) to cytokinesis, in (D). Line and error bars represent the mean and SD (N ≥ 50 cells from two independent experiments). The Mann–Whitney *U*-test (two-tailed) was used to obtain a *P*-value. \**P* < 0.0001. (**G**) Frequency of mitotic spindle structures upon CEP57 depletion. Values are presented as mean percentages ± SD (N = 6, triplicates, two independent experiments, at least 29 spindles in each assay).

### Pericentrin is crucial for bipolar spindle elongation in cells with two centrosomes

Although pericentrin and CDK5RAP2 are dispensable for efficient mitotic progression in cells with two centrosomes (Fig. 1A, B, 2C, S1G), the detailed functions of these PCM components in bipolar spindle formation have not been carefully examined. We subsequently analyzed the spindle length upon depletion of pericentrin or CDK5RAP2 in HeLa cells. We found that depletion of pericentrin significantly reduced the spindle length compared with that of control cells, whereas depletion of CDK5RAP2 had a limited effect on the spindle length (Fig. 6A, B). To further investigate this defect upon depletion of pericentrin, we performed live cell imaging of mitotic spindle formation in HeLa and HCT116 cells. Depletion of pericentrin delayed the elongation of two spindle poles (Fig. 6E, F, S4A, B). These results suggest that pericentrin supports spindle elongation. In line with this result, immunofluorescence analysis revealed that, in the pericentrin-depleted HeLa cells, the degree of CEP192 localization at the spindle poles was reduced; however, this was not observed in CDK5RAP2-depleted cells (Fig. 6C, D). This observation implies that pericentrin more efficiently recruits CEP192 to centrosomes, thereby facilitating spindle elongation.

**Figure 6.**
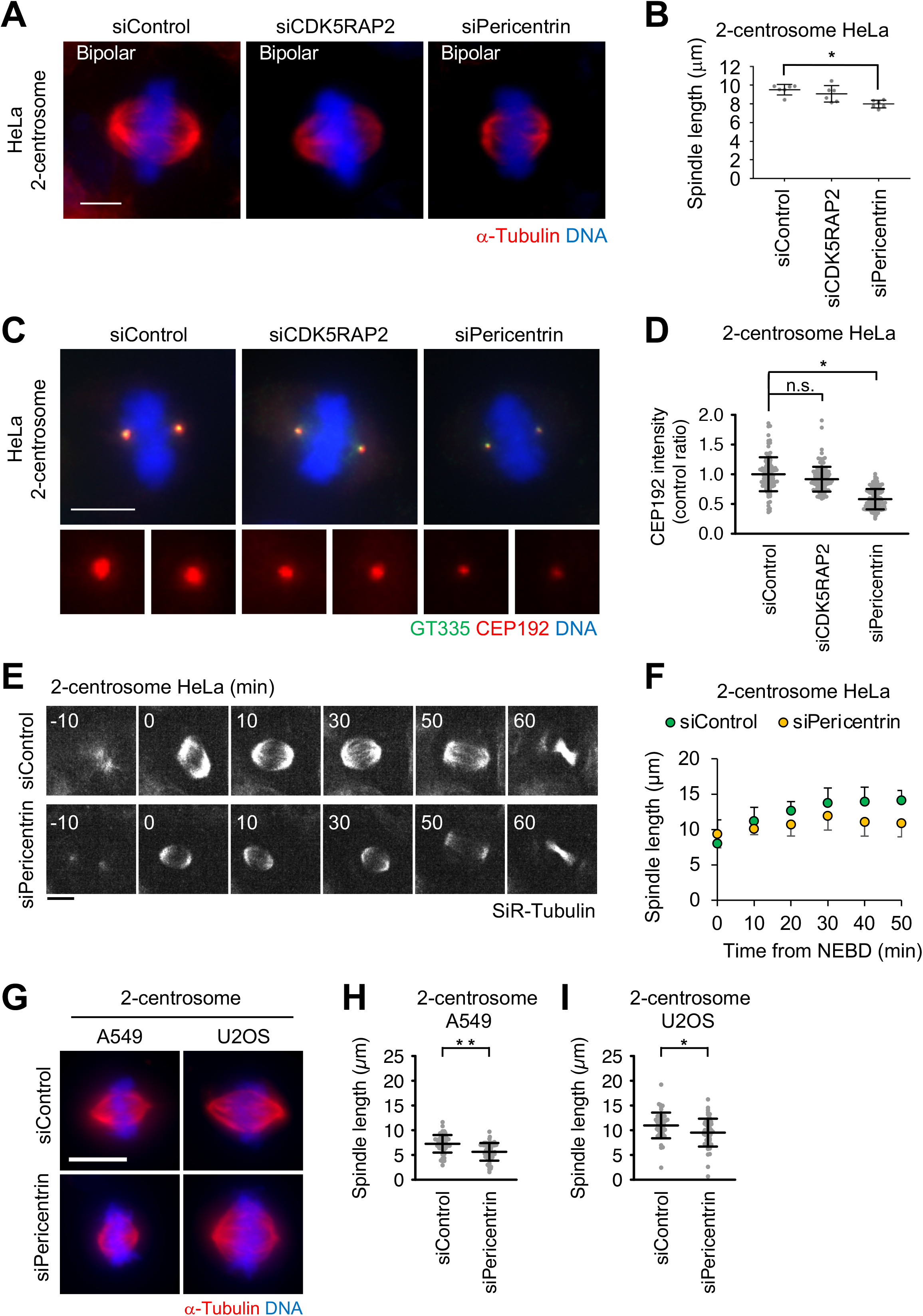
Pericentrin is crucial for the bipolar spindle elongation in cells with two centrosomes. (**A**) Mitotic spindle structures upon treatment with siRNA in cells with two centrosomes. Red and blue represent α-tubulin and DNA, respectively. Z-projections: 21 planes, 1 μm apart. Scale bar, 5 μm. (**B**) Quantification spindle length of HeLa cells (N > 14, triplicates, from two independent experiments). Line and error bars represent the mean and SD. One-way ANOVA with Tukey’s multiple comparisons test was used to determine the significance of the difference. \**P* < 0.005. (**C**) CEP192 observed in two-centrosome cells. Red, and blue represent GT335, CEP192, and DNA, respectively. Z-projections: 20 planes, 0.5 μm apart. Scale bar, 5 μm. (**D**) The signal intensity of CEP192 on centrosomes in (C). Line and error bars represent the mean and SD (N ≥ 50 cells from two independent experiments). Kruskal–Wallis test was used to determine the significance of the difference. \**P* < 0.0001. n.s., not significantly different. (**E**) Time-lapse observation of the structure of microtubules upon depletion of pericentrin in HeLa cells were observed with a 40× objective. Gray represent SiR-tubulin, respectively. Z-projections: 10 planes, 2.2 μm apart. Scale bar, 10 μm. Time zero corresponds to nuclear envelope break down (NEBD). (**F**) Averaged time courses of the pole length at each time point in (E). The length between two poles of spindle was measured from 40 cells from two independent experiments. Time course data were aligned at the time of the NEBD (0 min). Error bars, SD. (**G**) Mitotic spindle structures of A549 and U2OS cells. Red, and blue represent α-tubulin, and DNA, respectively. Z-projections: 31 planes, 0.5 μm apart. Scale bar, 10 μm. Quantification spindle length of A549 (H) and U2OS (I) cells (N > 40 from two independent experiments). Line and error bars represent the mean and SD. The Mann– Whitney *U*-test (two-tailed) was used to obtain a *P*-value. \**P* < 0.005, \*\**P* < 0.0001.

Furthermore, we tested the effect of depletion of pericentrin on the spindle elongation of various cell types. The spindle length in pericentrin-depleted A549, U2OS, A431, and PANC1 cells was significantly shorter than that noted in control cells (Fig. 6G, S4C, F–H). On the other hand, in some cell types (RPE1, GI1, SKOV3), the spindle length upon depletion of pericentrin was not altered compared with that observed in control cells (Fig. S4C–F). These results suggest that pericentrin is required for efficient spindle elongation in certain cell lines with two centrosomes.

### The activity of PLK1 is crucial for PCM-pole assembly and bipolar spindle formation in one-centrosome cells

The accumulation of PCM components at centrosomes in mitosis is regulated by PLK1 activity(Haren et al., 2009; Lee and Rhee, 2011; Joukov et al., 2014). However, we found that PLK1 and phosphorylated PLK1 were not detected at most PCM-poles in one-centrosome cells (Fig. 7A–D). To determine if PLK1 was required for PCM-pole assembly and subsequent bipolar spindle formation, we treated cells with a low dose of the PLK1 inhibitor BI 2536 (1 nM) and observed the amount of pericentrin at the centriolar pole and the spindle structure. Treatment of two-centrosome cells with a low dose of the PLK1 inhibitor caused chromosome congression errors and a slight reduction of pericentrin at centrosomes, but did not affect bipolar spindle formation (Fig. 7E–G). In contrast, in one-centrosome cells, PLK1 inhibition prevented PCM-pole formation and led to the formation of monopolar spindles (Fig. 7E, F). In addition, PLK1 inhibition greatly reduced the level of pericentrin at the centriolar pole compared with the level recorded in two-centrosome cells (Fig. 7G). Together, these results suggest that PLK1 activity is crucial for PCM-pole assembly and subsequent bipolar spindle formation in one-centrosome cells.

**Figure 7.**
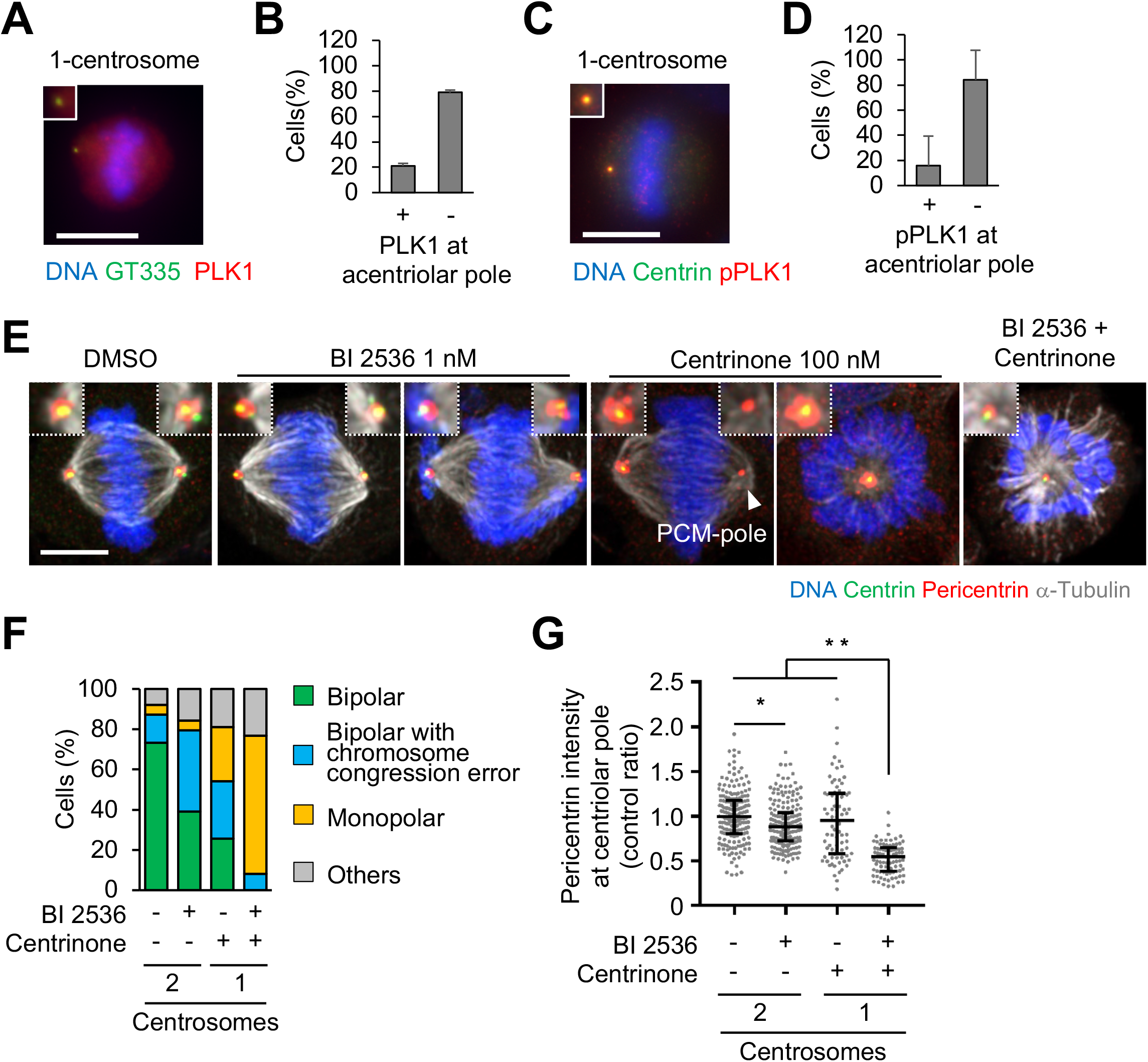
PLK1 is crucial for PCM-pole formation and bipolar spindle formation in one-centrosome cells. (**A, B**) PLK1 and phosphorylated PLK1 observed in one-centrosome cells. (**A**) Red, green, and blue represent PLK1, GT335, and DNA, respectively. Z-projections: 20 planes, 1 μm apart. Scale bar, 10 μm. (**B**) Frequency of localization of PLK1 in (A). Values are presented as mean percentages from two independent experiments (N > 40 from two experiments). (**C**) Red, green, and blue represent phosphorylated PLK1, centrin, and DNA, respectively. Z-projections of 20 sections, every 1 μm. Scale bar, 5 μm. (**D**) Frequency of localization of phosphorylated PLK1 in (A). Values are presented as mean percentages from triplicates (N > 40 from two experiments). (**E**) Mitotic spindle structures upon PLK1 inhibition with or without 100 nM of centrinone. HeLa cells expressing EGFP-centrin1 and Pericentrin-mCherry were observed with a 63× objective. Green, red, gray, and blue represent GFP (centrin1), RFP (Pericentrin), α-tubulin, and DNA, respectively. Z-projections: 10 planes, 0.3 μm apart. Scale bar, 5 μm. (**F**) Frequency of mitotic spindle structures in (A). Values are mean percentages from two independent experiments (N = 50 for each experiment). (**G**) The signal intensity of Pericentrin on centrin foci of fixed mitotic HeLa cells expressing EGFP-centrin1 and Pericentrin-mCherry (N > 45 for each condition). Line and error bars represent median with interquartile range. Kruskal–Wallis test was used to determine the significance of the difference. \**P* < 0.05, \*\**P* < 0.0001.

### Dual inhibition of PLK1 and PLK4 prevents cell growth in a wide variety of cancer cell lines

Mitotic spindle formation is a common target of anti-cancer drugs(Dumontet and Jordan, 2010; Tischer and Gergely, 2019; Henriques et al., 2019). Since the dual inhibition of PLK1 and PLK4 (PLK1+4i) efficiently prevented bipolar spindle formation in one-centrosome cells (Fig. 7E, F), we further tested the potential of PLK1+4i as an anticancer strategy. PLK1+4i efficiently prevented HeLa cell growth (Fig. 8A). The half maximal inhibitory concentration (IC_50_) value of the PLK1 inhibitor against HeLa cells was 1.1 nM (Table S2). Using both PLK1 and PLK4 inhibitors decreased the IC_50_ value to 0.6 nM (Table S2). Therefore, inhibition of both PLK1 and PLK4 has an additive effect on the growth suppression of HeLa cells. Under these conditions, after depletion of centrosomes, most of HeLa cells started to die due to prolonged mitosis (Fig. 6B, C, S6). The results suggest that the toxicity of PLK1+4i may be caused by inhibition of PCM-pole formation. Furthermore, PLK1+4i-treated cells showed cleavage of the apoptosis marker poly(ADP-ribose)-polymerase 1 (PARP1), suggesting that this drug combination induces apoptosis in mitosis (Fig. 6D). In addition, we assessed the effect of PLK1+4i in 19 cancer cell lines. The PLK1 inhibitor suppressed the proliferation of cancer cell lines to different extents (IC_50_ values in Table S1). U2OS, K562, and HMV-II cells showed approximately 10-fold higher resistance against the PLK1 inhibitor compared with HeLa cells. However, PLK1+4i efficiently prevented the growth of various cancer cell lines, including PLK1 inhibitor-resistant cell lines such as U2OS and K562 (15 cell lines: IC_50_[−centrinone B]/IC_50_[+centrinone B] ≥ 1.5) (Fig. 6E and Table 1). Overall, these results suggest that dual inhibition of PLK1 and PLK4 would be an effective drug target against cancer proliferation.

**Figure 8.**
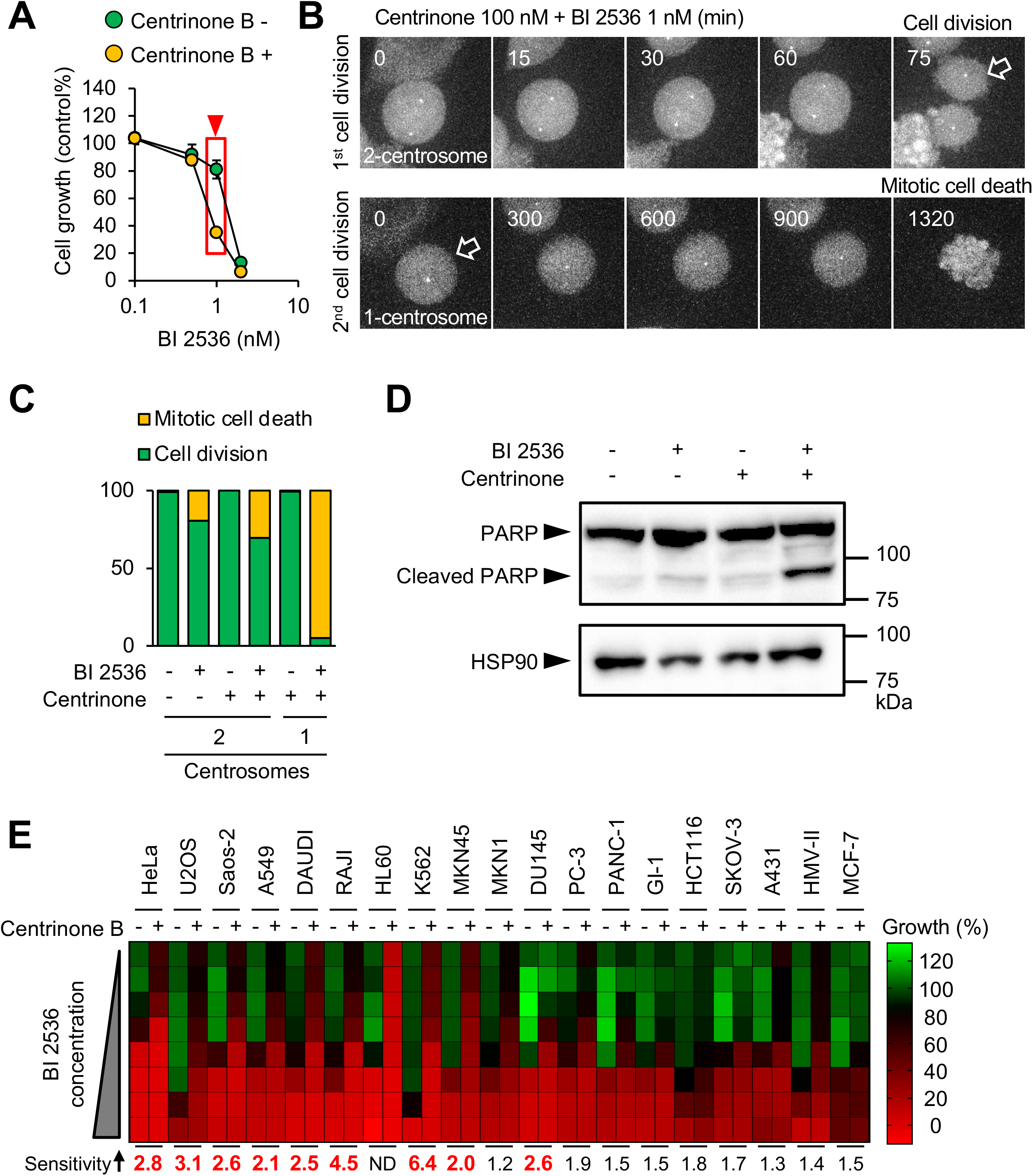
Dual inhibition of PLK1 and PLK4 prevents cancer cell proliferation. (**A**) Dual inhibition of PLK1 and PLK4 induced cell death in HeLa cells. Cell viability (% of DMSO or CentB mono-treatment) was determined after treatment with CentB and various concentrations of PLK1 inhibitor. (**B**) Mitotic cell fate of individual cells upon PLK1 and PLK4 dual inhibition. After centrosome depletion, treatment with a low dose of PLK1 inhibitor induced cell death. The mitotic cell fate of HeLa cells expressing EGFP-centrin1 and pericentrin-mCherry after treatment with 100 nM of centrinone, 1 nM of BI2536, or 100 nM of centrinone + 1 nM of BI2536 was observed compared to DMSO alone (solvent control). Cells were observed with a 60× objective. Z-projections: 30 planes, 1 μm apart. Scale bar, 5 μm. Time zero corresponds to the beginning of mitotic cell rounding. (**C**) Frequency of mitotic events in (B). Values are presented as percentages. N = 205 (for DMSO control, two centrosomes), 211 (for BI, two centrosomes), 107 (for centrinone, two centrosomes), 92 (for centrinone + BI, two centrosomes), 205 (for centrinone, one centrosome), 60 (for centrinone + BI, one centrosome) cells; data from two independent experiments were pooled. (**D**) Dual inhibition of PLK1 and PLK4 induced the cleavage of PARP. HeLa cells were treated with the drugs for 48 h and PARP cleavage was analyzed by western blotting. The concentrations of centrinone and BI 2536 were 100 nM and 1 nM, respectively. (**E**) Dual inhibition of both PLK1 and PLK4 in various cancer cell lines. After four days of treatment with 500 nM of centrinone B and various concentrations of BI2536 (0, 0.1, 0.5, 1, 2, 5, 10, or 20 nM), cell viability (% of DMSO control) was determined and is shown as a heat map. The ratios between IC_50_ values (± centrinone B) are shown below the heat map. Exact IC_50_ values are provided in Table 1.

## Discussion

In this study, we show that the centriole and PCM cooperate to recruit CEP192 at the spindle pole to facilitate bipolar spindle formation in human cells. We found that, even in cells in which PCM assembly was suppressed, CEP192 at the centriole wall efficiently promoted bipolar spindle assembly (Fig. 1). Furthermore, cells with one centrosome formed a bipolar spindle with a PCM-pole, which accumulates PCM proteins (including CEP192) at the opposite side of the centriolar spindle pole (Fig. 2–3). Consistently, the PCM-pole assembly is critical for cell division in one-centrosome cells (Fig. 4–5). Overall, the findings in this study illustrate that the centriole and PCM cooperatively promote bipolar spindle assembly through recruitment of CEP192 to the spindle pole in human somatic cells (Fig. 9). In addition, based on this evidence, we propose that dual inhibition of centriole duplication and a critical mitotic kinase PLK1 would be an attractive target for anti-cancer strategies (Fig. 7–8).

**Figure 9.**
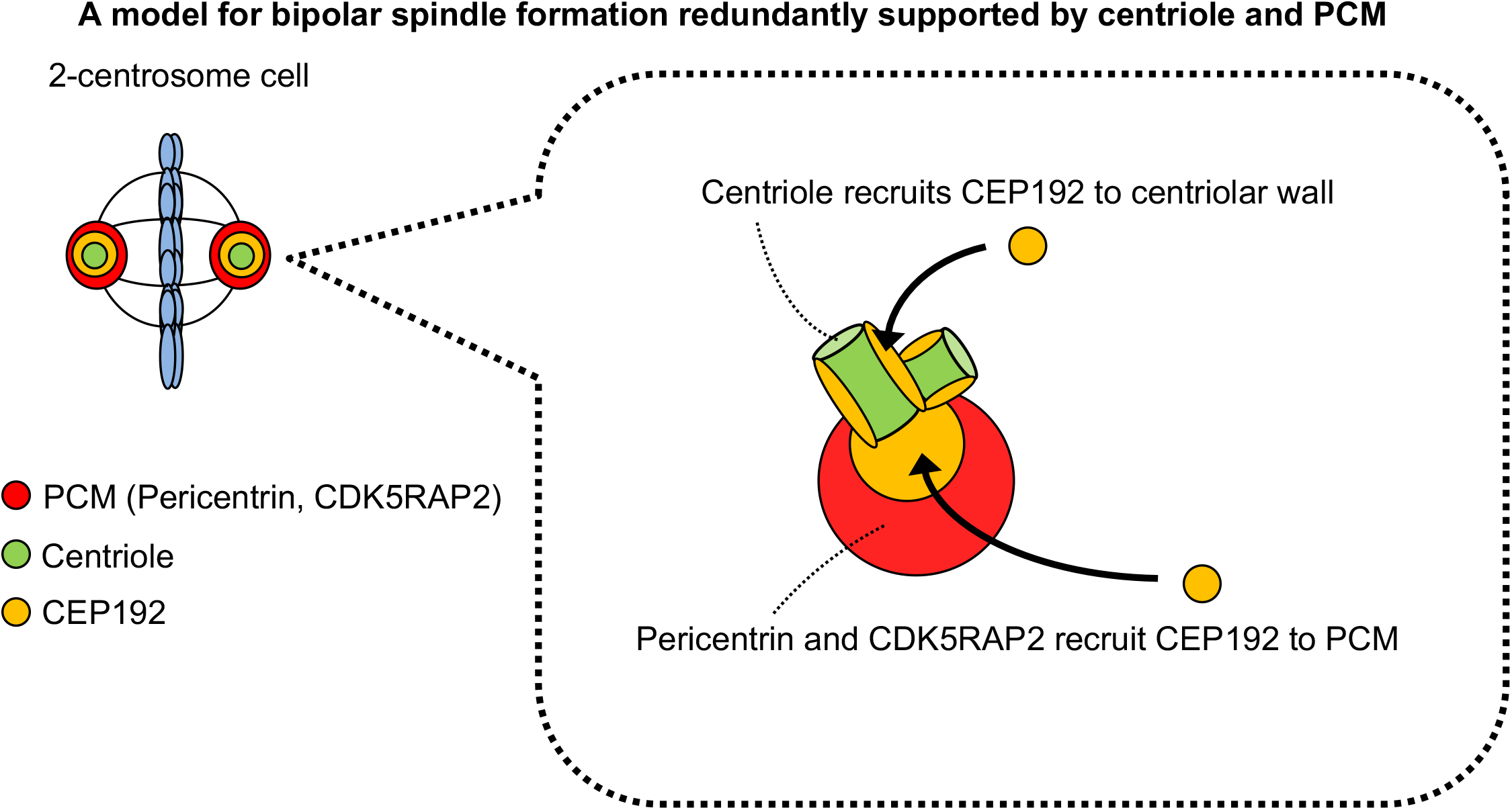
The Centriole and PCM cooperate to recruit CEP192 to the spindle pole to facilitate bipolar spindle formation. Schematic illustration of the assembly of CEP192 at the spindle pole by the centriole and PCM in human cells. For details, see the Discussion section.

In interphase cells, CEP192 localizes at the centrioles and regulates the microtubule nucleation activity of centrosomes(O’Rourke et al., 2014). In the G2/M phase, CEP192 is further recruited to PCM clouds by pericentrin(Joukov et al., 2014), promoting mitotic spindle formation. In pericentrin/CDK5RAP2 double-depleted cells, although the CEP192 localization was restricted on the centriolar wall, these cells efficiently completed mitosis (Fig. 1C–F). These results suggest that a fraction of CEP192 at the centriolar wall is sufficient for its function in mitosis. A previous study suggested that CEP192 supports the sequential activation of PLK1 and aurora kinase A (AURKA) at centrosomes(Joukov et al., 2014). Moreover, it has been shown that phosphorylated AURKA interacts with TPX2 and promotes spindle assembly(Joukov and De Nicolo, 2018). It is therefore possible that CEP192 at the centriole wall sufficiently activates the PLK1-AURKA pathway, thereby facilitating bipolar spindle formation.

We found that one-centrosome cells efficiently assembled PCM-poles (Fig. 2F, G, S2A-D). On the other hand, intriguingly, most zero-centrosome cells failed to assemble PCM proteins at the acentriolar poles (Chinen et al., 2020). How does this difference occur? In one-centrosome cells, PLK1 was localized only at centriolar poles, but not at PCM-poles (Fig. 7A–D). However, the PLK1 kinase activity is somehow necessary for the assembly of PCM-poles. It is possible that phosphorylation events at the centriole driven by the activity of PLK1 may provide a pool of PCM for the generation of the PCM-pole. In contrast, zero-centrosome cells do not have the platform components (e.g., centrioles) for PCM assembly. Previous research indicated that, in zero-centrosome cells, the activity of PLK1 in the cytoplasm was significantly increased(Takeda et al., 2020). However, in such cells, the PCM-pole was not assembled at the spindle poles(Takeda et al., 2020). Together, these observations suggest that the centriole itself is important for PCM assembly in human cells.

Knockdown experiments further revealed that CDK5RAP2 and pericentrin are crucial for cell division in one-centrosome cells. In addition, depletion of CEP57 augmented the assembly of PCM-poles, and facilitated mitotic progression in one-centrosome cells. These results indicate that the PCM proteins are required for PCM-pole formation in one-centrosome cells, and also raised the possibility that the balance of PCM quantities between two spindle poles may be a critical factor for proper mitotic progression in human cells. In line with this notion, it has been shown that in primary human malignancies, centrosome abnormalities such as centriole rosettes are frequently observed(Cosenza et al., 2017). These extra centrioles could lead to a greater accumulation of PCM proteins at the one centrosome, thereby increasing the nucleation of microtubules at this spindle pole and resulting in chromosome missegregation and aneuploidy. Our assay system may be useful for analyzing the balance of PCM quantities and the resulting microtubule nucleation between two spindle poles.

Based on the vulnerabilities of one-centrosome cells described above, our study also highlights the potential of dual inhibition of centriole duplication and PCM assembly as an attractive drug target for cancer therapies. The PLK1 inhibitor efficiently suppressed both PCM maturation and subsequent PCM-pole formation in one-centrosome cells. In this way, a low dose of PLK1 inhibitor efficiently suppressed cell division in one-centrosome cells, but not in two-centrosome cells (Fig. 8B, C). Previous clinical trials of PLK1 inhibitors have not been successful. Therefore, several studies have been performed to improve PLK1 inhibitor toxicity through combination with other inhibitors, such as α/β-tubulin inhibitors(Stehle et al., 2015; Weiß et al., 2015). In this study, the dual inhibition strategy, which inhibits both PLK1 and PLK4, provided an alternative approach to targeting PLK1 in the development of anticancer drugs. Interestingly, treatment with centrinone did not strongly alter the toxicity of microtubule inhibitors (Table S2). This result suggests that the strong toxicity caused by the dual inhibition of PLK1 and PLK4 was not merely due to an additive effect in mitosis, but rather the specific inhibition of both centrosomal and acentrosomal spindle assembly machinery. In addition, recently, it was suggested that decreased centrosome numbers are associated with poorer response to chemotherapy and an increased invasive capacity of tumor cells in ovarian cancer(Morretton et al., 2019). Therefore, our strategy to suppress PCM assembly in centrosome-reduced cells may be an attractive method for targeting ovarian cancer cells that have a reduced number of centrosomes.

## Supporting information

Fig1A_siCEP192

Fig1A_siControl

Fig1A_siPericentrin_siCDK5RAP2

Fig3B_pattern1

Fig3B_pattern2

Fig4C_siCDK5RAP2

Fig4C_siControl

Fig4C_siPericentrin

Fig5D_siCEP57

Fig5D_siControl

Fig6E_HeLa_siControl

Fig6E_HeLa_siPericentrin

Fig8B_1st cell_division_(2centrosome)

Fig8B_2nd cell_division_(1centrosome)

FigS1G_2centrosome_siCDK5RAP2

FigS1G_2centrosome_siCEP192

FigS1G_2centrosome_siControl

FigS1G_2centrosome_siPericentrin

FigS1H_1centrosome_siCDK5RAP2

FigS1H_1centrosome_siCEP192

FigS1H_1centrosome_siControl

FigS1H_1centrosome_siPericentrin

FigS1I_0centrosome_siCDK5RAP2

FigS1I_0centrosome_siCEP192

FigS1I_0centrosome_siControl

FigS1I_0centrosome_siPericentrin

FigS4A_HCT116_siControl

FigS4A_HCT116_siPericentrin

## Acknowledgements

We thank the Kitagawa lab members for fruitful discussions. We thank Dr. Andrew Shiau and Dr. Karen Oegema at Ludwig Institute for Cancer Research for providing centrinone B. We are also thankful to Dr. Y. Nagumo for providing the SKOV-3 cells. This work was supported by JSPS KAKENHI grants (Grant numbers: 24687026, 19H05651, 16H06168, 18K14705, and 17J02833) from the Ministry of Education, Science, Sports and Culture of Japan, Takeda Science Foundation, Mochida Memorial foundation, and Daiichi Sankyo Foundation of Life Science.

## Author contributions

T.C. and D.K. designed the study; T.C., K.Y., K.F., K.W., Y.T., and Y.N. performed the experiments; and T.C., K.Y., K.H., S.Y., Y.T., and D.K. designed the experiments. T.C., K.Y., Y.T., and D.T. analyzed the data; T.C., K.Y., and D.K. wrote the manuscript, which was reviewed by all authors.

## Methods

### Cell culture and transfection

HeLa and U2OS cells were obtained from the ECACC (European Collection of Authenticated Cell Cultures). These cell lines were authenticated by Short Tandem Repeat (STR) profiling at the ECACC. HeLa cells stably expressing EGFP-centrin1 have been previously described(Tsuchiya et al., 2016). HeLa cells expressing pericentrin endogenously tagged with mCherry were generated using the CRISPR/Cas9 system, as previously described, with slight modifications(Natsume et al., 2016). GuideRNA oligos (Pericentrin_gRNA_F: CACCGCTGTTTAATCATCGGGTGGC and Pericentrin_gRNA_R: AAACGCCACCCGATGATTAAACAGC) were hybridized and cloned into the BbsI site of pX330 (Addgene). To construct the donor plasmid for homology-directed repair, the homology arms of the *Pericentrin* locus (chr21:47864730-47865813) were amplified (pBS2_Pericentrin C-ter_InsF: GGTATCGATAAGCTTACCAGGTAATGCAAGTCCTCGCCG and pBS2_Pericentrin C-ter_InsR: CGCTCTAGAACTAGTAGAATGCTCCGGGTTCCACTGA) from the genomic DNA of HeLa cells and cloned into pBluescript using the Infusion Cloning kit (Takara). A BamHI sequence with a silent mutation to prevent re-cutting was generated in the middle of the homology arm domain by mutagenesis PCR (Pericentrin C-ter silent BamHI_F: TACTTCAAAGAAATCTTGCCACCCGATGATTAAACAGGGATCCATAAAATGTCATGGCTCTTTCCTGCGA, Pericentrin C-ter silent BamHI_R: GCCATGACATTTTATGGATCCCTGTTTAATCATCGGGTGGCAAGATTTCTTTGAAGTAGAATCTGCATATAAATAAAAATGAGG). The mCherry cassette containing a hygromycin-resistant gene was introduced into the BamHI site in the middle of the homology arms. The plasmids were introduced into the HeLa cell line stably expressing EGFP-centrin1(Tsuchiya et al., 2016) and isolated using the limited dilution method with hygromycin. Saos-2, A549, DAUDI, RAJI, HL60, K562, MKN45, MKN1, DU145, PC-3, PANC-1, GI-1, A431, HMV-II, and MCF-7 were obtained from the RIKEN BioResource Research Center. These cell lines were authenticated by STR profiling at the RIKEN BioResource Research Center. HCT116 cells were obtained from the American Type Culture Collection (ATCC CCL-247). HCT116 CMVOsTIR1 HsSAS6–AID have been previously described(Yoshiba et al., 2019). HCT116 cell lines were cultured in McCoy’s 5A medium (Thermo Fisher Scientific) supplemented with 10% fetal bovine serum, 2 mM L-glutamine, 100 U/ml penicillin, and 100 μg/ml streptomycin. SKOV-3 was provided Dr. Yoko Nagumo. HeLa, U2OS, A549, GI-1, and A431 cells were cultured in Dulbecco’s modified Eagle’s medium containing 10% fetal bovine serum, 100 U/ml penicillin and 100 μg/ml streptomycin at 37 °C in a 5% CO_2_ atmosphere. Saos-2, DAUDI, RAJI, HL60, MKN45, MKN1, DU145, PC-3, PANC-1, and SKOV-3 cells were cultured in RPMI1680 medium containing 10% fetal bovine serum, 100 U/ml penicillin, and 100 μg/ml streptomycin at 37 °C in a 5% CO_2_ atmosphere. K562 and HMV-II cells were cultured in Ham’s F12 medium containing 10% fetal bovine serum, 100 U/ml penicillin, and 100 μg/ml streptomycin at 37 °C in a 5% CO_2_ atmosphere. MCF-7 cells were cultured in MEM medium containing 10% fetal bovine serum, 1 mM sodium pyruvate, 100 U/ml penicillin, and 100 μg/ml streptomycin at 37 °C in a 5% CO_2_ atmosphere. Transfection of siRNA constructs was conducted using Lipofectamine RNAiMAX (Life Technologies). Unless otherwise noted, the transfected cells were analyzed 48 h after transfection with siRNA.

### RNA interference

The following siRNAs were used: Silencer Select siRNA (Life Technologies) against CEP57 (s18692), CEP192 (s226819), CDK5RAP2 (s31429, s31430), pericentrin (s10136, s10138), and negative control #1 (4390843).

### Chemicals

The following chemicals were used in this study: Centrinone (PLK4 inhibitor, MedChem Express, HY-18682), Centrinone B (PLK4 inhibitor, gift from Dr. Andrew Shiau and Dr. Karen Oegema), BI2536 (PLK1 inhibitor, A10134; AdooQ), proTAME (APC/C inhibitor, I-440, Boston Biochem), Cytochalasin B (actin inhibitor, Wako, 036-17553), paclitaxel (α/β-tubulin stabilizer: Wako, 163-18614), SiR-Tubulin (Microtubule probe, CY-SC002; SPIROCHROME).

### Antibodies

The following primary antibodies were used in this study: rabbit polyclonal antibodies against CDK5RAP2 (IHC00063: immunofluorescence [IF] 1:500, Western Blotting [WB] 1:1000; Bethyl Laboratories), Cep192 (A302–324A, IF 1:1,000, WB 1:1000; Bethyl Laboratories), Cep152 (A302–480A, IF 1:1,000; Bethyl Laboratories), CP110 (12780–1-AP, IF 1:1,000; Proteintech), CPAP/CENP-J (11517-1-AP, IF 1:100; Proteintech), and α-tubulin (PM054, IF 1:300; MBL); mouse monoclonal antibodies against γ-tubulin (GTU88) (T6557, IF 1:1,000; Sigma–Aldrich), Polyglutamylation Modification (GT335, mAb) (AG-20B-0020-C100, IF 1:1,000; AdipoGen), α-tubulin (T5168, IF 1:1,000; Sigma-Aldrich), HSP90 (610419, WB 1:5000; BD Biosciences) pericentrin (ab4448, WB 1:1,000; Abcam) and pericentrin (ab28144, IF 1:1,000; Abcam). Fluorescein isothiocyanate-labeled anti-GFP was purchased from Abcam (ab6662, IF 1:250 or 1:500). Alexa 488-labeled Cep152 (A302–480A, IF 1:250; Bethyl Laboratories) was generated with Alexa Fluor labeling kits (Life Technologies) and used for three color staining. The following secondary antibodies were used: Alexa Fluor 488 goat anti-mouse IgG (H+L) (A-11001, 1:500; Molecular Probes), Alexa Fluor 555 goat anti-rabbit IgG (H+L) (A-21428, 1:500; Molecular Probes), and Alexa Fluor 647 goat anti-mouse IgG (ab150115, 1:500; Abcam).

### Sample preparations for immunostaining

Cells were treated with 100 nM of centrinone or 500 nM of centrinone B for 1–3 days to induce acentrosomal cells (Fig. 2A, B, F, G, S2A, B). To observe the microtubule nucleation from PCM-poles (Fig. S2E), firstly, HeLa cells expressing pericentrin-mCherry and EGFP-centrin1 were treated with 100 nM of centrinone for two days. Then, cells were arrested in metaphase through treatment with 20 μM of proTAME for 4 h and incubated on ice for 1 h to depolymerize microtubules. Subsequently, cells were incubated at 25 °C for 5 min. For the Sas-6 depletion experiments using the AID system (Fig. S2C, D), the cells were incubated with 50 μM of indole-3-acetic acid (IAA) for two days. To observe CEP192 localization at acentriolar poles (Fig. 4G), cells were treated with 100 nM of centrinone and siRNA for two days. Then, cells were arrested in metaphase through treatment with 20 μM of proTAME for 6 h. To examine the effect of siRNA on acentriolar cells (Fig. 4A, B, 5A, B, G, S3), cells were treated with 100 nM of centrinone or 500 nM of centrinone B and siRNA for two days. For the chemical perturbation experiments (Fig. 7E–G, 8D), cells were treated with 100 nM of centrinone and BI 2536 for two days.

### Western blotting

For preparation of total cell lysates, cells were lysed in 1× SDS sample buffer. SDS– PAGE was performed using 6 or 10% polyacrylamide gels, followed by transfer on Immobilon-P membrane (Millipore Corporation). Blocking was performed in 2.5% skim milk in PBS containing 0.02% Tween (PBS-T) for 30 min at room temperature. The membrane was probed with the primary antibodies for 12-18 h at 4°C, washed with PBS-T three times. After that, the membrane was incubated with HRP-conjugated secondary antibodies for 1 h at room temperature and washed with PBS-T three times. The signals were detected with ECL Prime/Select reagents (GE Healthcare) or Chemi-Lumi One Ultra (Nacalai Tesque) via the ChemiDoc XRSþ system (Bio-Rad).

### Sample preparations for live-cell imaging

For live cell imaging, HeLa cells stably expressing EGFP-centrin1, HeLa cells expressing pericentrin-mCherry and EGFP-centrin1, HeLa cells expressing CDK5RAP2-mCherry and EGFP-centrin1, HeLa cells expressing mCherry–NuMA and EGFP–centrin1 were cultured in 35-mm glass-bottom dishes (#627870; Greiner Bio-One) or 24-well SENSOPLATE (#662892; Greiner Bio-One) at 37 °C in a 5% CO_2_ atmosphere.

To observe the dynamics of PCM (Fig. 3A–E) in one-centrosome or zero-centrosome cells, cells were treated with 500 nM of centrinone B. To test the effect of depletion of PCM proteins or CEP57 on one-centrosome cells (Fig. 4C–F, 5D–F), cells were treated with siRNA with 100 nM of centrinone for two days. To test the effect of drug combinations on mitotic cell fate (Fig. 8B, C, S5), the cell cycle progression of HeLa cells expressing pericentrin-mCherry and EGFP-centrin1 was observed in the presence of 100 nM centrinone and 1 nM BI 2536. To simultaneously observe one-centrosome cells and zero-centrosome cells (Fig. S2C-E, S1G-I), after two days of treatment with 0.1% DMSO or 100 nM centrinone (to enrich zero-centrosome cells), HeLa cells expressing EGFP-centrin1 were treated with siRNA with 0.1% DMSO or 100 nM of centrinone. Prior to imaging, cells were incubated with 100 nM of SiR-Tubulin for 4 h to visualize the microtubules.

### Microscopy for immunofluorescence analyses

For immunofluorescence analyses, the cells cultured on coverslips (No. 1; Matsunami) were fixed using methanol at −20 °C for 7 min and washed with PBS. The cells were permeabilized after fixation with PBS/0.05% TritonX-100 (PBSX) for 5 min, and blocked in 1% BSA in PBSX for 30 min at room temperature. The cells were then incubated with primary antibodies for 7–24 h at 4 °C, washed thrice with PBSX, and incubated with secondary antibodies and 0.2 μg ml^−1^ Hoechst 33258 (DOJINDO) for 45–60 min at room temperature. The cells were washed thrice with PBSX and mounted onto glass slides.

We counted the number of spindle patterns using a DeltaVision Personal DV-SoftWoRx system (Applied Precision) or an Axioplan2 fluorescence microscope (Carl Zeiss). Confocal microscopy images were captured by the Leica TCS SP8 system. For deconvolution for confocal microcopy images, Huygens essential software (Scientific Volume Imaging) was used.

STED images were taken using a Leica TCS SP8 STED 3X system with a Leica HCPL APO 100 × /1.40 oil STED WHITE and a 660 nm laser line for depletion. Scan speed was set to 100 Hz with 5× line averaging in a 512 × 80 px format (pixel size 15– 20 nm). The Z interval was set to 180 nm. The STED images were processed by deconvolution using the Huygens Professional software (SVI).

Maximum intensity z-projections of a representative picture for each condition were generated using the FIJI distribution of the ImageJ software. The number and step sizes of z-planes are described in the figure legends.

### Microscopy for live imaging

A Confocal Scanner Box, Cell Voyager CV1000 (Yokogawa Electric Corp.) equipped with a 60 × oil immersion objective or CQ1 Benchtop High-Content Analysis System equipped with a 40× objective was used for live cell imaging. Imaging was initiated after the addition of drugs or 24–48 h after transfection, and images were acquired every 10 min for 24–48 h. Maximum intensity z-projections of representative images for each condition were generated using the FIJI distribution of the ImageJ software. The number and step sizes of z-planes are described in the figure legends.

### Cell viability

Cell viability was determined using the WST-8 assay. Exponentially growing cells (1 × 10^3^ cells/well in a 96-well plate) were treated with compounds for three (Fig. 8A, Table S2) or four (Fig. 8E, Table S1) days. WST-8 assay reagents (DOJINDO) were added to the culture. After several hours of incubation, the absorbance at 450 nm was measured with FilterMax F3 & F5 Multi-Mode Microplate Readers (Molecular Devices). After subtracting the background (blank well), cell viability (control %) was determined.

### Statistical analysis

Statistical analyses were performed using the GraphPad Prism 7 software. Except for Fig. 6B, p-values were determined by non-parametric methods (Mann–Whitney U-test or Kruskal–Wallis test). P-value in Fig. 6B was determined by One-way ANOVA with Tukey’s multiple comparisons test Details are described in the figure legends.

### Data availability

The data supporting the findings of this study are available from the corresponding authors upon request.

---

**Supplementary Figure 1.**
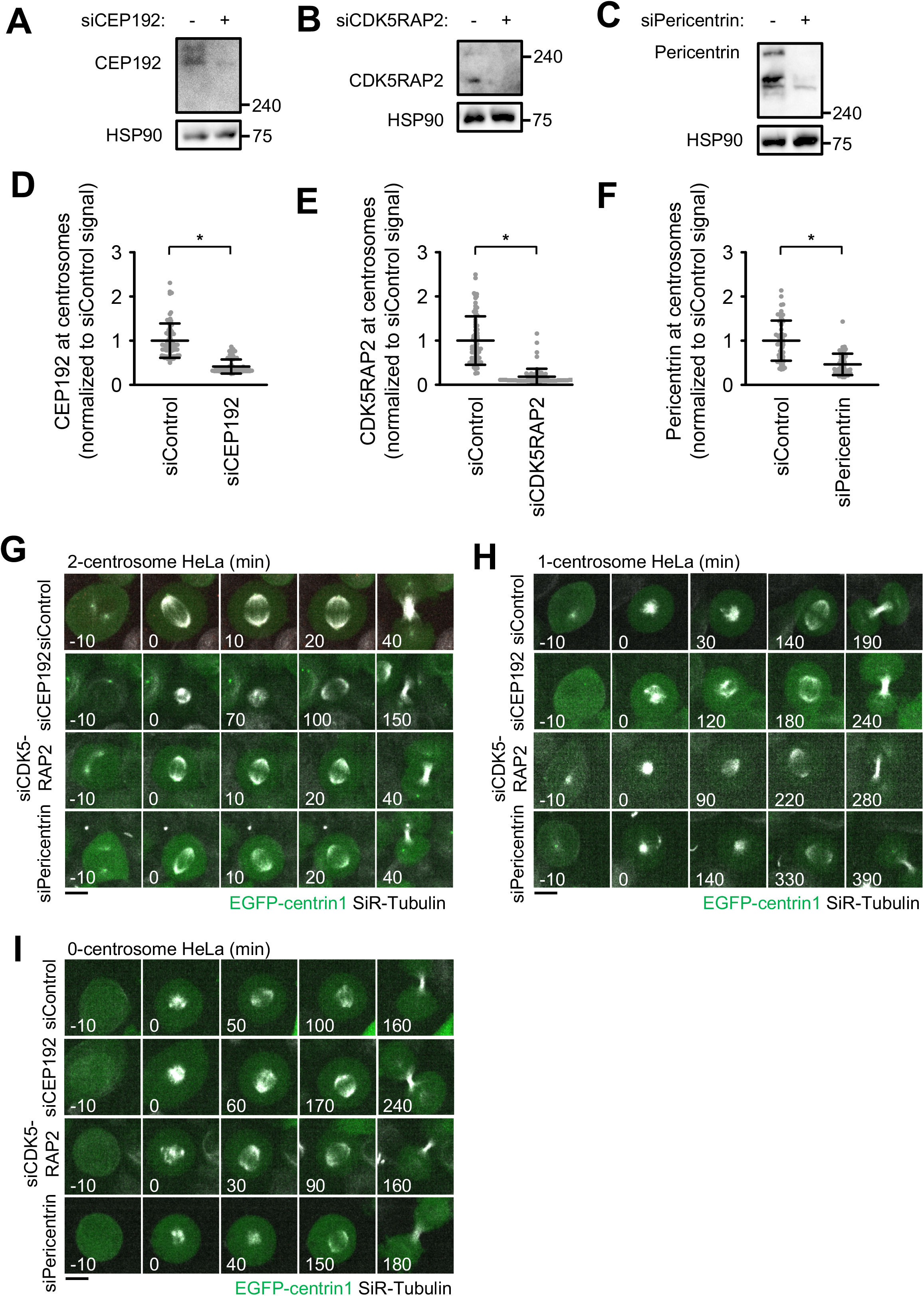
Depletion of PCM components in two-, one-, and zero-centrosome cells. (A-C) Western blot analysis of the efficiency of protein depletion of CEP192 (A), CDK5RAP2 (B) and pericentrin (C) after 48 h of siRNA transfection in HeLa cells. (**D-F**) Quantification of depleted centrosomal CEP192 (D), CDK5RAP2 (E) and pericentrin The Mann–Whitney *U*-test (two-tailed) was used to obtain a *P*-value. (*P* < 0.0001). (**G-I**). Time-lapse observation of the structure of microtubules upon siRNA treatment against the indicated proteins. DMSO-treated two-centrosome (G), centrinone-treated one-centrosome (H), and centrinone-treated zero-centrosome (I) HeLa cells expressing EGFP-centrin1 and pericentrin-mCherry were observed with a 40× objective. Gray represent SiR-tubulin, respectively. Z-projections: 10 planes, 2.2 μm apart. Scale bar, 10μm. Time zero corresponds to nuclear envelope break down (NEBD).

**Supplementary Figure 2.**
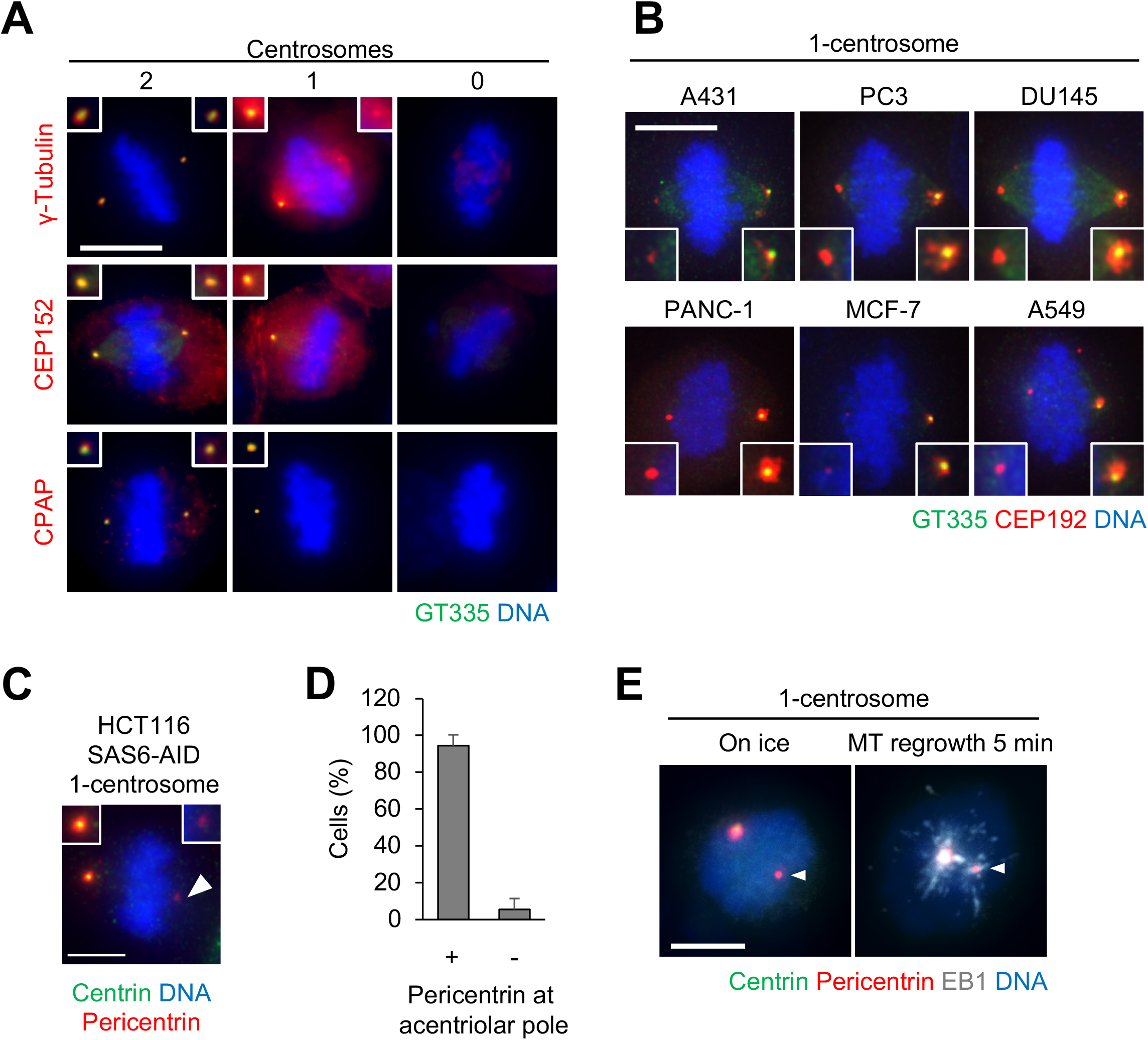
Distribution of centrosomal factors in centriolar and acentriolar spindle poles. (**A–D**) Distribution of centrosomal factors in centriolar and acentriolar spindle poles. (**A**) DMSO-treated control mitotic spindles (two centrosomes) and centrinone-treated spindles (one or zero centrosomes) of HeLa cells. Green, red, and blue represent GT335, protein of interest (γ-tubulin, CEP152, or CPAP), and DNA, respectively. Z-projections: 21 planes, 1 μm apart. Scale bar, 10 μm. (**B**) PCM-poles were observed in various cells. Magenta, cyan, and blue represent CEP192, GT335, and DNA, respectively. Z-projections: 40 planes, 0.3 μm apart. Scale bar, 5 μm. (**C**) PCM-poles were observed in one-centrosome spindles induced by SAS6 depletion. Red, green, and blue represent centrin, pericentrin, and DNA, respectively. Scale bar, 5 μm. (**D**) Quantification of pole patterns in (C). Values are presented as mean percentages from three independent experiments (N = 20 for each experiment). (**E**) Microtubule nucleation from the PCM-pole. Following treatment with ice, microtubule nucleation (5 min at 25°C) was observed in one-centrosome cells. Gray, red, green, and blue in the merged image represent EB1, pericentrin, centrin, and DNA, respectively. Z-projections: 21 planes, 1 μm apart. Scale bar, 5 μm.

**Supplementary Figure 3.**
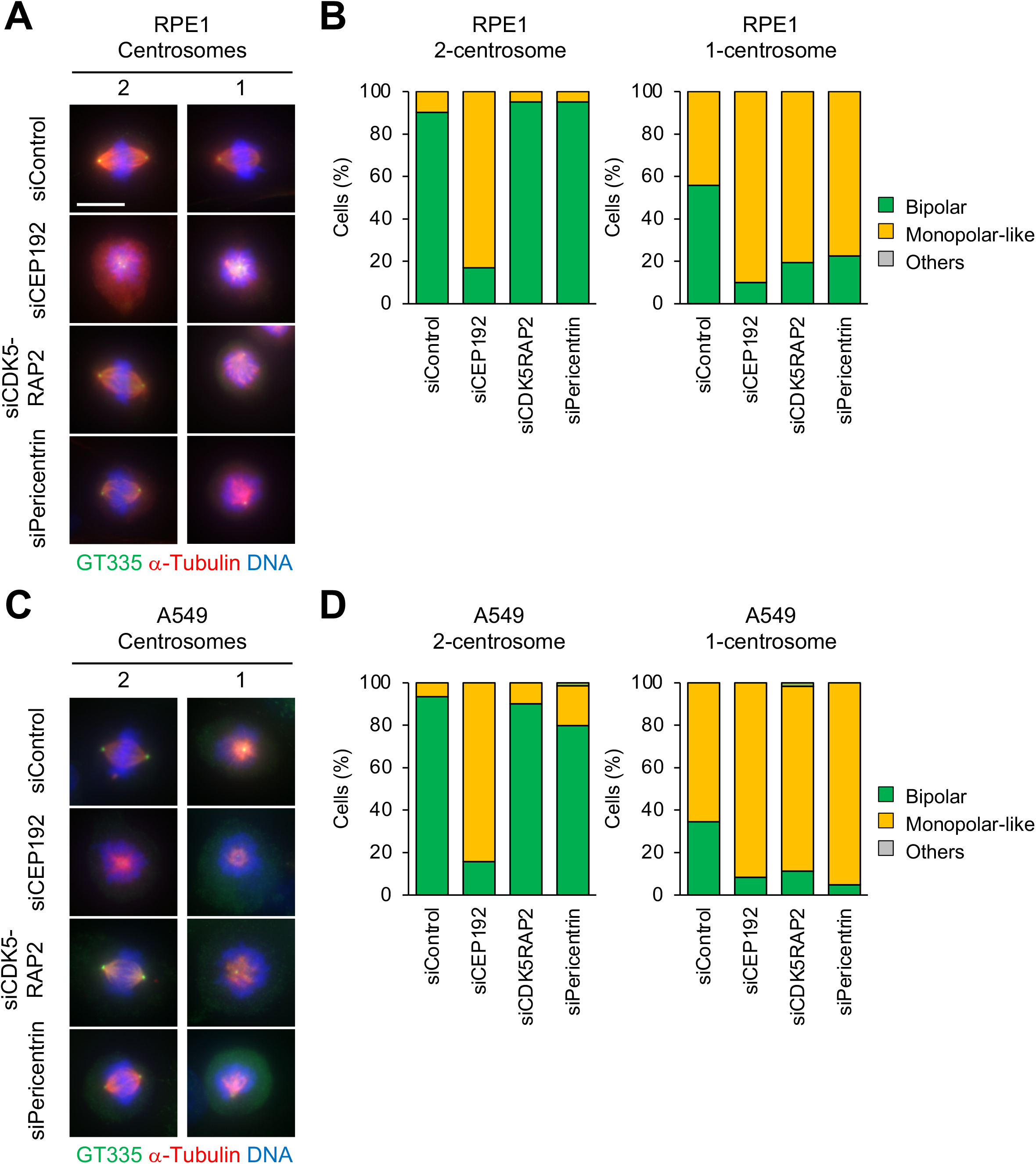
Pericentrin and CDK5RAP2 are crucial for the cell division of one-centrosome cells. (**A-D**) Mitotic spindle structures in siPCM-treated RPE1 (A) and A549 (C) cells. After a 2-day siRNA treatment with or without 100 nM of centrinone, mitotic spindle structures were observed with a 63× objective. Green, red, and blue represent GT335, α-tubulin, and DNA, respectively. Z projections: 31 planes, 0.5 μm apart. Scale bar, 10 μm. Frequency of mitotic spindle structures in RPE1 (B) and A549 (D) cells. Values are mean percentages from two independent experiments (N > 20 for each experiment).

**Supplementary Figure 4.**
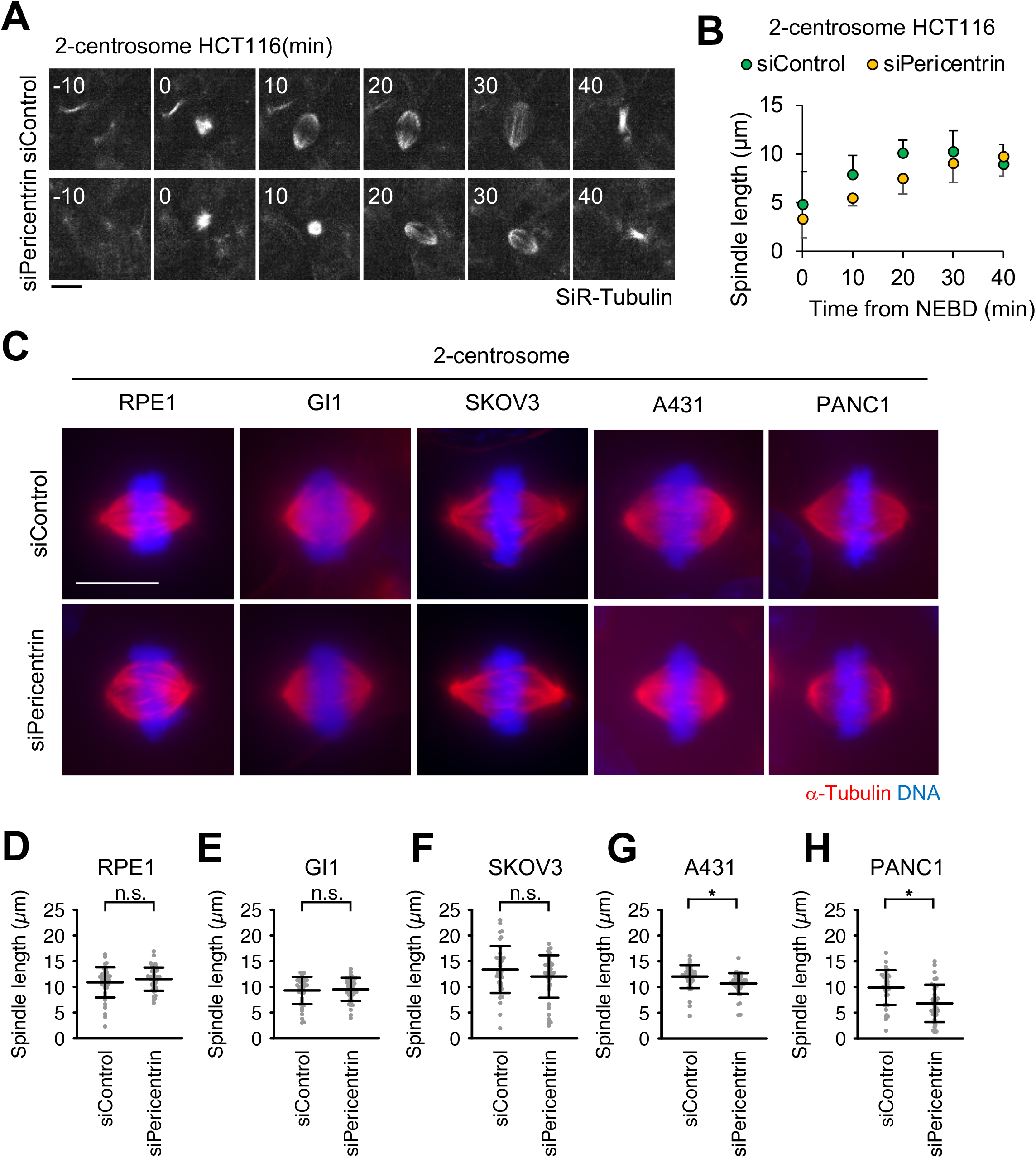
Pericentrin is critical for proper spindle elongation in two-centrosome cells. (**A**) Time-lapse observation of the structure of microtubules upon siRNA treatment against the indicated proteins. HCT116 cells were observed with a 40× objective. Gray represent SiR-tubulin, respectively. Z-projections: 10 planes, 2.2 μm apart. Scale bar, 10 μm. Time zero corresponds to nuclear envelope break down (NEBD). (**B**) Averaged time courses of the pole length at each time point in (A). The length between two poles of spindle was measured from 40 cells from two independent experiments. Time course data were aligned at the time of the NEBD (0 min). Error bars, SD. (**C**) The time required for the establishment of two spindle poles in (A). Line and error bars represent the mean and SD (N ≥ 75 cells from two independent experiments). The Mann–Whitney *U*-test (two tailed) was used to obtain a *P*-value. (*P* < 0.05). (**D**) Mitotic spindle structures of RPE1, GI1, SKOV3, A431 and PANC1 cells. Red, and blue represent α-tubulin, and DNA, respectively. Z-projections: 31 planes, 0.5 μm apart. Scale bar, 10 μm. (**E**-**I**) Quantification spindle length (N > 40 from two independent experiments) in (D). Line and error bars represent the mean and SD. The Mann–Whitney *U*-test (two-tailed) was used to obtain a *P*-value. (*P* < 0.0005). n.s., not significantly different.

**Supplementary Figure 5.**
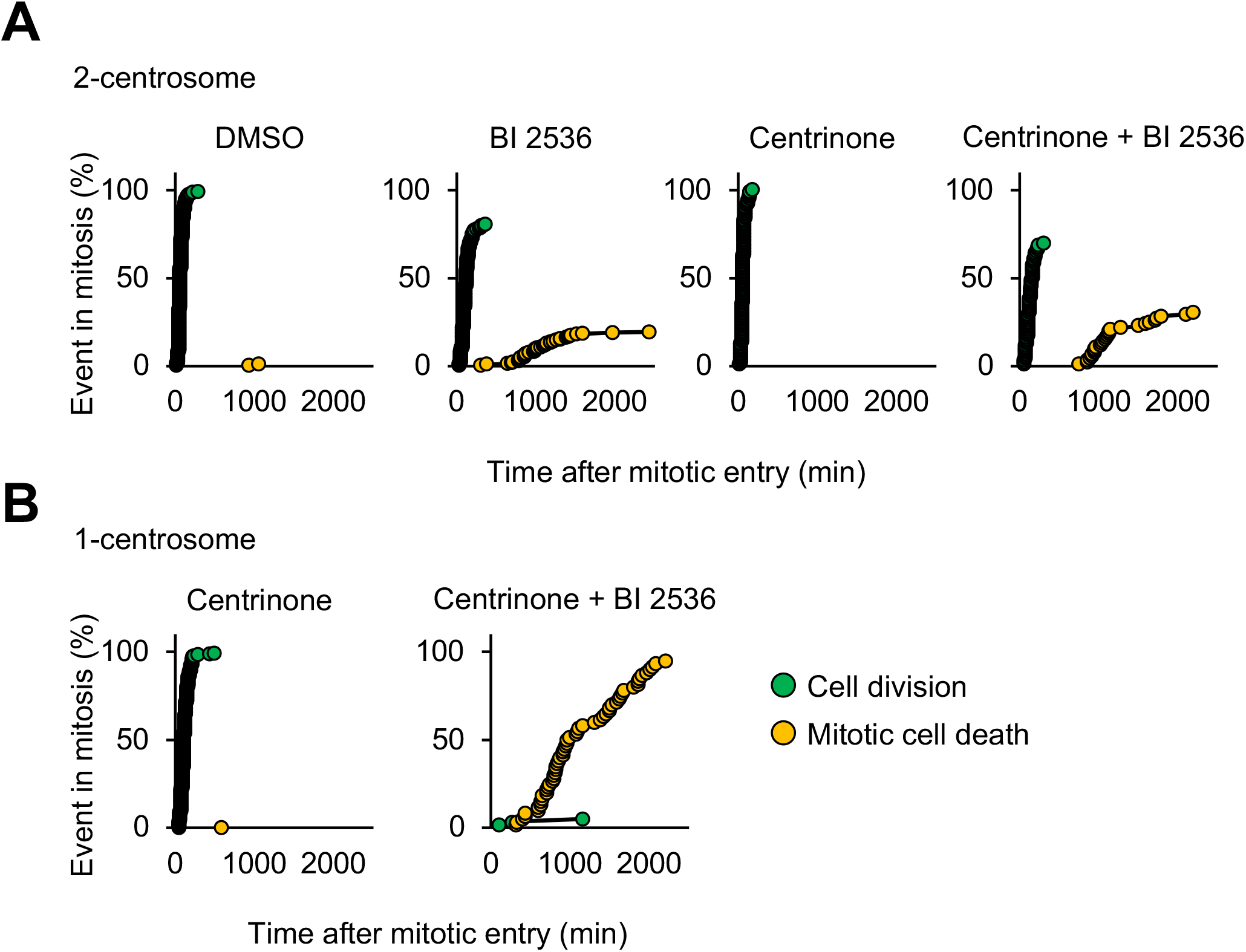
Dual inhibition of PLK1 and PLK4 prolongs mitosis and induces cell death in one-centrosome HeLa cells. Each plot shows the cumulative percentage of cell division or cell death in mitosis at each time point of Fig. 6B, C. N = 205 (for DMSO control, two centrosomes), 211 (for BI, two centrosomes), 107 (for centrinone, two centrosomes), 92 (for centrinone + BI, two centrosomes), 205 (for centrinone, one centrosome), 60 (for centrinone + BI, one centrosome) cells; data from two independent experiments were pooled.

**Table S1.** IC_50_ values of BI 2536 against the growth of various cancer cells with or without centrinone B (CentB). Various cancer cells were treated with BI 2536 with or without 500 nM of centrinone B for four days and their viabilities were determined using the WST-8 assay. IC_50_ values and the ratio between IC_50_ values (±CentB) are shown. Related to Fig. 6E.

| Cell line | Type of cancer | IC<sub>50</sub> values<br>of BI2536<br>(nM)<br>(–CentB) | IC<sub>50</sub> values<br>of BI2536<br>(nM)<br>(+CentB) | Ratio of IC<sub>50</sub> values<br>(–CentB/+CentB) |
| --- | --- | --- | --- | --- |
| HeLa | Cervical cancer | 1.1 ± 0.1 | 0.4 ± 0.2 | 2.8 |
| U2OS | Osteosarcoma | 11.0 ± 3.9 | 3.6 ± 0.6 | 3.1 |
| Saos-2 | Osteosarcoma | 2.1 ± 0.4 | 0.8 ± 0.3 | 2.6 |
| A549 | Lung carcinoma | 2.3 ± 0.6 | 1.1 ± 0.2 | 2.1 |
| DAUDI | Burkitt lymphoma | 3.5 ± 0.6 | 1.4 ± 0.3 | 2.5 |
| RAJI | Burkitt lymphoma | 4.5 ± 0.9 | 1.0 ± 0.1 | 4.5 |
| HL60 | Leukemia | Not<br>determined | Not<br>determined | Not determined |
| K562 | Leukemia | 13.4 ± 0.5 | 2.1 ± 0.6 | 6.4 |
| MKN45 | Gastric cancer | 4.7 ± 1.3 | 2.4 ± 0.3 | 2.0 |
| MKN1 | Gastric cancer | 4.1 ± 0.9 | 3.4 ± 1.3 | 1.2 |
| DU145 | Prostate carcinoma brain<br>metastasis | 3.1 ± 0.7 | 1.2 ± 0.4 | 2.6 |
| PC-3 | Prostate carcinoma bone<br>metastasis | 3.7 ± 1.8 | 2.0 ± 0.9 | 1.9 |
| PANC-1 | Pancreatic carcinoma | 3.3 ± 0.8 | 2.2 ± 0.6 | 1.5 |
| GI-1 | Glioma | 3.5 ± 0.2 | 2.4 ± 0.4 | 1.5 |
| HCT116 | Colon cancer | 5.7 ± 4.0 | 3.2 ± 1.6 | 1.8 |
| SKOV-3 | Ovarian cancer | 2.9 ± 0.6 | 1.7 ± 0.3 | 1.7 |
| A431 | Epidermoid carcinoma | 2.5 ± 0.5 | 2.0 ± 0.9 | 1.3 |
| HMV-II | Melanoma | 10.6 ± 3.0 | 7.6 ± 4.3 | 1.4 |
| MCF-7 | Breast adenocarcinoma | 6.4 ± 1.6 | 4.3 ± 1.3 | 1.5 |

**Table S2.** IC_50_ values of mitotic inhibitors against the growth of HeLa cells with or without centrinone B (CentB). HeLa cells were treated with several compounds with or without 500 nM of centrinone B for 3 days and their viabilities were determined by WST-8 assay. IC_50_ values (average from at least two independent assays) and the ratio between IC_50_ values (±CentB) were shown.

| Compounds (Targets) | IC<sub>50</sub> values (- CentB) | IC<sub>50</sub> values (+CentB) | Ratio of IC<sub>50</sub> values (- CentB/+CentB) |
| --- | --- | --- | --- |
| BI2536 (PLK1) | 1.1 nM | 0.6 nM | 1.8 |
| Cytochalasin B (β-actin) | 1.5 μM | 1.6 μM | 0.9 |
| Paclitaxel ($\alpha/\beta$-tubulin) | 1.8 nM | 1.7 nM | 1.1 |

## References

Alvarez-Rodrigo, I., T.L. Steinacker, S. Saurya, P.T. Conduit, J. Baumbach, Z.A. Novak, M.G. Aydogan, A. Wainman, and J.W. Raff. 2019. Evidence that a positive feedback loop drives centrosome maturation in fly embryos. Elife. 8:1–31. doi:10.7554/eLife.50130.

Baumann, C., X. Wang, L. Yang, and M.M. Viveiros. 2017. Error-prone meiotic division and subfertility in mice with oocyte-conditional knockdown of pericentrin. J. Cell Sci. 130:1251–1262. doi:10.1242/jcs.196188.

Bettencourt-Dias, M., A. Rodrigues-Martins, L. Carpenter, M. Riparbelli, L. Lehmann, M.K. Gatt, N. Carmo, F. Balloux, G. Callaini, and D.M. Glover. 2005. SAK/PLK4 is required for centriole duplication and flagella development. Curr. Biol. 15:2199– 2207. doi:10.1016/j.cub.2005.11.042.

Cabral, G., T. Laos, J. Dumont, and A. Dammermann. 2019. Differential Requirements for Centrioles in Mitotic Centrosome Growth and Maintenance. Dev. Cell. 50:355–366.e6. doi:10.1016/j.devcel.2019.06.004.

Chavali, P.L., G. Chandrasekaran, A.R. Barr, P. Tátrai, C. Taylor, E.K. Papachristou, C.G. Woods, S. Chavali, and F. Gergely. 2016. A CEP215–HSET complex links centrosomes with spindle poles and drives centrosome clustering in cancer. Nat. Commun. 7:11005. doi:10.1038/ncomms11005.

Chen, C.-T., H. Hehnly, Q. Yu, D. Farkas, G. Zheng, S.D. Redick, H.-F. Hung, R. Samtani, A. Jurczyk, S. Akbarian, C. Wise, A. Jackson, M. Bober, Y. Guo, C. Lo, and S. Doxsey. 2014. A Unique Set of Centrosome Proteins Requires Pericentrin for Spindle-Pole Localization and Spindle Orientation. Curr. Biol. 24:2327–2334. doi:10.1016/j.cub.2014.08.029.

Chinen, T., S. Yamamoto, Y. Takeda, K. Watanabe, K. Kuroki, K. Hashimoto, D. Takao, and D. Kitagawa. 2020. NuMA assemblies organize microtubule asters to establish spindle bipolarity in acentrosomal human cells. EMBO J. 39:e102378. doi:10.15252/embj.2019102378.

Choi, Y., P. Liu, S.K. Sze, C. Dai, and R.Z. Qi. 2010. CDK5RAP2 stimulates microtubule nucleation by the γ-tubulin ring complex. J. Cell Biol. 191:1089– 1095. doi:10.1083/jcb.201007030.

Clift, D., and M. Schuh. 2015. A three-step MTOC fragmentation mechanism facilitates bipolar spindle assembly in mouse oocytes. Nat. Commun. 6:7217. doi:10.1038/ncomms8217.

Conduit, P.T., K. Brunk, J. Dobbelaere, C.I. Dix, E.P. Lucas, and J.W. Raff. 2010. Centrioles Regulate Centrosome Size by Controlling the Rate of Cnn Incorporation into the PCM. Curr. Biol. 20:2178–2186. doi:10.1016/j.cub.2010.11.011.

Conduit, P.T., J.H. Richens, A. Wainman, J. Holder, C.C. Vicente, M.B. Pratt, C.I. Dix, Z.A. Novak, I.M. Dobbie, L. Schermelleh, and J.W. Raff. 2014. A molecular mechanism of mitotic centrosome assembly in Drosophila. 1–23. doi:10.7554/eLife.03399.

Consolati, T., J. Locke, J. Roostalu, Z.A. Chen, J. Gannon, J. Asthana, W.M. Lim, F. Martino, M.A. Cvetkovic, J. Rappsilber, A. Costa, and T. Surrey. 2020. Microtubule Nucleation Properties of Single Human γTuRCs Explained by Their Cryo-EM Structure. Dev. Cell. 53:603–617.e8. doi:10.1016/j.devcel.2020.04.019.

Cosenza, M.R., A. Cazzola, A. Rossberg, N.L. Schieber, G. Konotop, E. Bausch, A. Slynko, T. Holland-Letz, M.S. Raab, T. Dubash, H. Glimm, S. Poppelreuther, C. Herold-Mende, Y. Schwab, and A. Krämer. 2017. Asymmetric Centriole Numbers at Spindle Poles Cause Chromosome Missegregation in Cancer. Cell Rep. 20:1906–1920. doi:10.1016/j.celrep.2017.08.005.

Dudka, D., C. Castrogiovanni, N. Liaudet, H. Vassal, and P. Meraldi. 2019. Spindle-Length-Dependent HURP Localization Allows Centrosomes to Control Kinetochore-Fiber Plus-End Dynamics. Curr. Biol. 29:3563–3578.e6. doi:10.1016/j.cub.2019.08.061.

Dumontet, C., and M.A. Jordan. 2010. Microtubule-binding agents: a dynamic field of cancer therapeutics. Nat. Rev. Drug Discov. 9:790–803. doi:10.1038/nrd3253.

Erpf, A.C., L. Stenzel, N. Memar, M. Antoniolli, M. Osepashvili, R. Schnabel, B. Conradt, and T. Mikeladze-Dvali. 2019. PCMD-1 Organizes Centrosome Matrix Assembly in C. elegans. Curr. Biol. 29:1324–1336.e6. doi:10.1016/j.cub.2019.03.029.

Gomez-Ferreria, M.A., U. Rath, D.W. Buster, S.K. Chanda, J.S. Caldwell, D.R. Rines, and D.J. Sharp. 2007. Human Cep192 Is Required for Mitotic Centrosome and Spindle Assembly. Curr. Biol. 17:1960–1966. doi:10.1016/j.cub.2007.10.019.

Gönczy, P. 2015. Centrosomes and cancer: revisiting a long-standing relationship. Nat. Rev. Cancer. 15:639–652. doi:10.1038/nrc3995.

Habedanck, R., Y.D. Stierhof, C.J. Wilkinson, and E.A. Nigg. 2005. The Polo kinase Plk4 functions in centriole duplication. Nat. Cell Biol. 7:1140–1146. doi:10.1038/ncb1320.

Hanafusa, H., S. Kedashiro, M. Tezuka, M. Funatsu, S. Usami, F. Toyoshima, and K. Matsumoto. 2015. PLK1-dependent activation of LRRK1 regulates spindle orientation by phosphorylating CDK5RAP2. Nat. Cell Biol. 17:1024–1035. doi:10.1038/ncb3204.

Haren, L., T. Stearns, and J. Lüders. 2009. Plk1-Dependent Recruitment of γ-Tubulin Complexes to Mitotic Centrosomes Involves Multiple PCM Components. PLoS One. 4:e5976. doi:10.1371/journal.pone.0005976.

Henriques, A.C., D. Ribeiro, J. Pedrosa, B. Sarmento, P.M.A. Silva, and H. Bousbaa. 2019. Mitosis inhibitors in anticancer therapy: When blocking the exit becomes a solution. Cancer Lett. 440–441:64–81. doi:10.1016/j.canlet.2018.10.005.

Hyman, A.A. 2014. Pericentriolar material structure and dynamics. Philos. Trans. R. Soc. Lond. B. Biol. Sci. 369. doi:10.1098/rstb.2013.0459.

Joukov, V., and A. De Nicolo. 2018. Aurora-PLK1 cascades as key signaling modules in the regulation of mitosis. Sci. Signal. 11:eaar4195. doi:10.1126/scisignal.aar4195.

Joukov, V., J.C. Walter, and A. De Nicolo. 2014. The Cep192-Organized Aurora A-Plk1 Cascade Is Essential for Centrosome Cycle and Bipolar Spindle Assembly. Mol. Cell. 55:578–591. doi:10.1016/j.molcel.2014.06.016.

Kim, J., J. Kim, and K. Rhee. 2019. PCNT is critical for the association and conversion of centrioles to centrosomes during mitosis. J. Cell Sci. 132:jcs225789. doi:10.1242/jcs.225789.

Kim, J., K. Lee, and K. Rhee. 2015. PLK1 regulation of PCNT cleavage ensures fidelity of centriole separation during mitotic exit. Nat. Commun. 6:10076. doi:10.1038/ncomms10076.

Kim, S., and K. Rhee. 2014. Importance of the CEP215-Pericentrin Interaction for Centrosome Maturation during Mitosis. PLoS One. 9:e87016. doi:10.1371/journal.pone.0087016.

Kirkham, M., T. Müller-Reichert, K. Oegema, S. Grill, and A.A. Hyman. 2003. SAS-4 Is a C. elegans Centriolar Protein that Controls Centrosome Size. Cell. 112:575– 587. doi:10.1016/S0092-8674(03)00117-X.

Kollman, J.M., A. Merdes, L. Mourey, and D.A. Agard. 2011. Microtubule nucleation by γ-tubulin complexes. Nat. Rev. Mol. Cell Biol. 12:709–721. doi:10.1038/nrm3209.

Lecland, N., and J. Lüders. 2014. The dynamics of microtubule minus ends in the human mitotic spindle. Nat. Cell Biol. 16:770–778. doi:10.1038/ncb2996.

Lee, K., and K. Rhee. 2011. PLK1 phosphorylation of pericentrin initiates centrosome maturation at the onset of mitosis. J. Cell Biol. 195:1093–1101. doi:10.1083/jcb.201106093.

Lee, S., and K. Rhee. 2010. CEP215 is involved in the dynein-dependent accumulation of pericentriolar matrix proteins for spindle pole formation. Cell Cycle. 9:775–784. doi:10.4161/cc.9.4.10667.

Liu, P., E. Zupa, A. Neuner, A. Böhler, J. Loerke, D. Flemming, T. Ruppert, T. Rudack, C. Peter, C. Spahn, O.J. Gruss, S. Pfeffer, and E. Schiebel. 2020. Insights into the assembly and activation of the microtubule nucleator γ-TuRC. Nature. 578:467– 471. doi:10.1038/s41586-019-1896-6.

Moritz, M., M.B. Braunfeld, J.W. Sedat, B. Alberts, and D.A. Agard. 1995. Microtubule nucleation by γ-tubulin-containing rings in the centrosome. Nature. 378:638–640. doi:10.1038/378638a0.

Morretton, J.-P., A. Herbette, C. Cosson, B. Mboup, A. Latouche, P. Gestraud, T. Popova, M.-H. Stern, F. Nemati, D. Decaudin, G. Bataillon, V. Becette, D. Meseure, A. Nicolas, O. Mariani, C. Bonneau, J. Barbazan, A. Vincent-Salomon, F. Mechta-Grigoriou, S.R. Roman, R. Rouzier, X. Sastre-Garau, O. Goundiam, and R. Basto. 2019. Centrosome amplification favours survival and impairs ovarian cancer progression. bioRxiv. 623983. doi:10.1101/623983.

Natsume, T., T. Kiyomitsu, Y. Saga, and M.T. Kanemaki. 2016. Rapid Protein Depletion in Human Cells by Auxin-Inducible Degron Tagging with Short Homology Donors. Cell Rep. 15:210–218. doi:10.1016/j.celrep.2016.03.001.

Nigg, E.A., and J.W. Raff. 2009. Centrioles, Centrosomes, and Cilia in Health and Disease. Cell. 139:663–678. doi:10.1016/j.cell.2009.10.036.

O’Rourke, B.P., M.A. Gomez-Ferreria, R.H. Berk, A.M.U. Hackl, M.P. Nicholas, S.C. O’Rourke, L. Pelletier, and D.J. Sharp. 2014. Cep192 controls the balance of centrosome and non-centrosomal microtubules during interphase. PLoS One. 9. doi:10.1371/journal.pone.0101001.

Schuh, M., and J. Ellenberg. 2007. Self-Organization of MTOCs Replaces Centrosome Function during Acentrosomal Spindle Assembly in Live Mouse Oocytes. Cell. 130:484–498. doi:10.1016/j.cell.2007.06.025.

Seo, M.Y., W. Jang, and K. Rhee. 2015. Integrity of the Pericentriolar Material Is Essential for Maintaining Centriole Association during M Phase. PLoS One. 10:e0138905. doi:10.1371/journal.pone.0138905.

Sir, J., M. Pütz, O. Daly, C.G. Morrison, M. Dunning, J. V Kilmartin, and F. Gergely. 2013. Loss of centrioles causes chromosomal instability in vertebrate somatic cells. J. Cell Biol. 203:747–756. doi:10.1083/jcb.201309038.

Stehle, A., M. Hugle, and S. Fulda. 2015. Eribulin synergizes with Polo-like kinase 1 inhibitors to induce apoptosis in rhabdomyosarcoma. Cancer Lett. 365:37–46. doi:10.1016/j.canlet.2015.04.011.

Takeda, Y., K. Yamazaki, K. Hashimoto, K. Watanabe, T. Chinen, and D. Kitagawa. 2020. The centriole protein CEP76 negatively regulates PLK1 activity in the cytoplasm for proper mitotic progression. J. Cell Sci. jcs.241281. doi:10.1242/jcs.241281.

Teixido-Travesa, N., J. Roig, and J. Luders. 2012. The where, when and how of microtubule nucleation - one ring to rule them all. J. Cell Sci. 125:4445–4456. doi:10.1242/jcs.106971.

Tischer, J., and F. Gergely. 2019. Anti-mitotic therapies in cancer. J. Cell Biol. 218:10– 11. doi:10.1083/jcb.201808077.

Tsuchiya, Y., S. Yoshiba, A. Gupta, K. Watanabe, and D. Kitagawa. 2016. Cep295 is a conserved scaffold protein required for generation of a bona fide mother centriole. Nat. Commun. 7:12567. doi:10.1038/ncomms12567.

Tungadi, E.A., A. Ito, T. Kiyomitsu, and G. Goshima. 2017. Human microcephaly ASPM protein is a spindle pole-focusing factor that functions redundantly with CDK5RAP2. J. Cell Sci. 130:3676–3684. doi:10.1242/jcs.203703.

Watanabe, K., D. Takao, K.K. Ito, M. Takahashi, and D. Kitagawa. 2019. The Cep57-pericentrin module organizes PCM expansion and centriole engagement. Nat. Commun. 10:931. doi:10.1038/s41467-019-08862-2.

Weiß, L.M., M. Hugle, S. Romero, and S. Fulda. 2015. Synergistic induction of apoptosis by a polo-like kinase 1 inhibitor and microtubule-interfering drugs in Ewing sarcoma cells. Int. J. Cancer. n/a-n/a. doi:10.1002/ijc.29725.

Wieczorek, M., L. Urnavicius, S.-C. Ti, K.R. Molloy, B.T. Chait, and T.M. Kapoor. 2020. Asymmetric Molecular Architecture of the Human γ-Tubulin Ring Complex. Cell. 180:165–175.e16. doi:10.1016/j.cell.2019.12.007.

Wong, Y.L., J. V Anzola, R.L. Davis, M. Yoon, A. Motamedi, A. Kroll, C.P. Seo, J.E. Hsia, S.K. Kim, J.W. Mitchell, B.J. Mitchell, A. Desai, T.C. Gahman, A.K. Shiau, and K. Oegema. 2015. Reversible centriole depletion with an inhibitor of Polo-like kinase 4. Science (80-.). 348:1155–1160. doi:10.1126/science.aaa5111.

Woodruff, J.B., B. Ferreira Gomes, P.O. Widlund, J. Mahamid, A. Honigmann, and A.A. Hyman. 2017. The Centrosome Is a Selective Condensate that Nucleates Microtubules by Concentrating Tubulin. Cell. 169:1066–1077.e10. doi:10.1016/j.cell.2017.05.028.

Woodruff, J.B., O. Wueseke, and A.A. Hyman. 2014. Pericentriolar material structure and dynamics. Phil. Trans. R. Soc. B. 369:20130459.

Yoshiba, S., Y. Tsuchiya, M. Ohta, A. Gupta, G. Shiratsuchi, Y. Nozaki, T. Ashikawa, T. Fujiwara, T. Natsume, M.T. Kanemaki, and D. Kitagawa. 2019. HsSAS-6-dependent cartwheel assembly ensures stabilization of centriole intermediates. J. Cell Sci. 132:jcs217521. doi:10.1242/jcs.217521.

Zheng, Y., M.L. Wong, B. Alberts, and T. Mitchison. 1995. Nucleation of microtubule assembly by a γ-tubulin-containing ring complex. Nature. 378:578–583. doi:10.1038/378578a0.

Zhu, F., S. Lawo, A. Bird, D. Pinchev, A. Ralph, C. Richter, T. Müller-Reichert, R. Kittler, A.A. Hyman, and L. Pelletier. 2008. The Mammalian SPD-2 Ortholog Cep192 Regulates Centrosome Biogenesis. Curr. Biol. 18:136–141. doi:10.1016/j.cub.2007.12.055.

